# Three-doses of BNT162b2 COVID-19 mRNA vaccine establishes long-lasting CD8^+^ T cell immunity in CLL and MDS patients

**DOI:** 10.1101/2022.05.13.491706

**Authors:** Susana Patricia Amaya Hernandez, Ditte Stampe Hersby, Kamilla Kjærgaard Munk, Tripti Tamhane, Darya Trubach, Maria Tagliamonte, Luigi Buonaguro, Anne Ortved Gang, Sine Reker Hadrup, Sunil Kumar Saini

## Abstract

Patients with hematological malignancies are prioritized for COVID-19 vaccine due to their high risk for severe SARS-CoV-2 infection related disease and mortality. To understand T cell immunity, its long-term persistence, and correlation with antibody response, we evaluated the BNT162b2 COVID-19 mRNA vaccine-specific immune response in chronic lymphocytic leukemia (CLL) and myeloid dysplastic syndrome (MDS) patients. Longitudinal analysis of CD8^+^ T cells using DNA-barcoded peptide-MHC multimers covering the full SARS-CoV-2 Spike-protein (415 peptides) showed vaccine-specific T cell activation and persistence of memory T cells up to six months post-vaccination. Surprisingly, a higher frequency of vaccine-induced antigen-specific CD8^+^ T cell was observed in the patient group compared to a healthy donor group. Furthermore, and importantly, immunization with the second booster dose significantly increased the frequency of antigen-specific CD8^+^ T cells as well as the total number of T cell specificities. Altogether 59 BNT162b2 vaccine-derived immunogenic epitopes were identified, of which 23 established long-term CD8^+^ T cell memory response with a strong immunodominance for NYNYLYRLF (HLA-A24:02) and YLQPRTFLL (HLA-A02:01) epitopes. In summary, we mapped the vaccine-induced antigen-specific CD8^+^ T cells and showed a booster-specific activation and enrichment of memory T cells that could be important for long-term disease protection in this patient group.

**Key Points:**

- COVID-19 mRNA vaccine induced an early and persistent activation of antigen-specific CD8^+^ T cells in this patient group.
- Vaccination with a booster dose is required to maintain vaccine-specific CD8^+^ T cells.

## Introduction

Patients with chronic lymphocytic leukemia (CLL) and Myelodysplastic syndrome (MDS) are generally faced with immune defects and deficiencies, related to their primary disease and as result of their cancer treatments, that may impact the immune cells’ ability to evade infections.^1–3^ Due to increased risk for severe SARS-CoV-2 infection and mortality,^4–6^ patients with hematological malignancies (HM) were prioritized for a vaccine-mediated protective immunity, initially with a two-dose COVID-19 vaccine and subsequently with an additional booster dose. However, we have limited knowledge on vaccine-induced T cell immunity, which could be critical for proper assessment of vaccine-mediated long-term protection.^7,8^ Especially since inconsistent and impaired vaccine-induced humoral response have been reported in HM patients.^9–13^ Here, we report antigen-specific CD8^+^ T cell response induced by BNT162b2 COVID-19 mRNA vaccine up to six-months post vaccination, and provide a detailed characterization of T cell immunodominance, immunological memory, and impact of the booster immunization in CLL and MDS patients.

## Study design

We used DNA-barcoded peptide-major histocompatibility complex (pMHC) multimers to evaluate CD8^+^ T cell response to the BNT162b2 mRNA vaccine, including the impact of booster immunization, in patients with pre-existing hematological cancers (CLL, n=23; MDS, n=5; Supplemental Table 1) up to six months post-vaccination and compared with a cohort of healthy individuals (n=19; Supplemental Table 2) (Figure 1A). By using NetMHCpan4.1^14^, we selected 415 unique peptides (8-11 amino acids) from the SARS-CoV-2 Spike protein encoded by the BNT162b2 mRNA vaccine (GenBank ID: QHD43416.1). The peptides were predicted to bind one or more of the nine prevalent HLA-A and HLA-B molecules generating 506 peptide-HLA pairs for experimental evaluation (Figure 1B, Supplemental Table 3). Additionally, 67 peptides from cytomegalovirus (CMV), Epstein-Barr viral (EBV), and influenza (FLU), (together denoted as CEF) were included to compare Spike to CEF-antigen-reactive T cells (Supplemental Table 4). DNA-barcoded multimers were prepared by loading HLA-specific peptides on MHC molecules and multimerized on dextramer backbone tagged with DNA-barcodes, providing each pMHC multimer with a unique DNA-barcode.^15^ PBMCs were incubated with HLA-matching (Supplemental Tables 5-6) pMHC multimers and stained with a phenotype antibody panel (Supplemental Table S7) to identify multimer-reactive CD8^+^ T cells (Supplemental Figure 1; see supplemental methods).^16^

**Figure 1.**
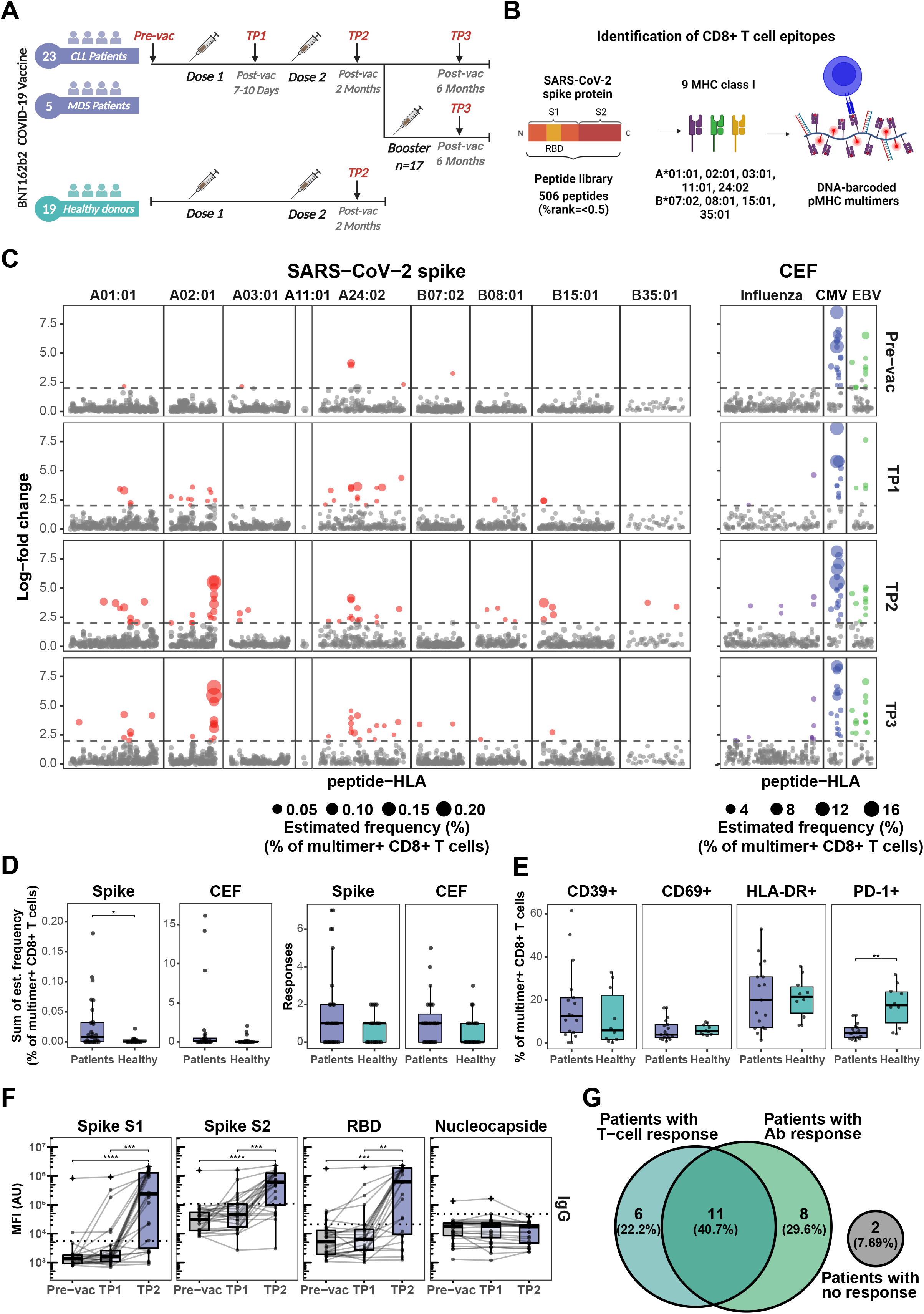
BNT162b2 COVID-19 mRNA vaccine drives early and persistent antigen-specific CD8+ T cells activation in CLL and MDS patients. (A) Sample collection timeline for patients with hematological malignancies (CLL, n=23 individuals; MDS, n=5 individuals) and healthy donors’ cohort (n=19 individuals). All donors received either two or three (booster) doses of the BNT162b2 mRNA vaccine. (B) Representation of the peptide library and the DNA-barcoded peptide-MHC (pMHC) multimers used to analyze T cell recognition towards the SARS-CoV-2 Spike protein derived HLA-binding peptides. (C) Summary of CD8^+^ T cell recognition to SARS-CoV-2 Spike- and CEF-derived peptides in the HM patients at four time points: prior to vaccination (Pre-vac), 7 to 10 days after first vaccine dose (TP1), 2 months after first vaccination dose (TP2), and 6 months after first vaccine dose (TP3). Responses were identified based on the enrichment of DNA barcodes associated with each of the tested peptide specificities (LogFc ≥ 2 and p < 0.001, Barracoda). Each dot represents one peptide-HLA combination per patient. The size of each significant response (colored dotes) is proportional to the estimated frequency (%) calculated from the percentage read count of the associated barcode out of the percentage of CD8^+^ multimer^+^ T cells. SARS-CoV-2-specific T cell responses were segregated based on their HLA. (D) Box plots comparing the number of significant SARS-CoV-2 Spike- and CEF-derived peptides responses (right) and the sum of their estimated frequencies (left) for each individual in the HM patients and the healthy donors at TP2. Each dot represents one sample. Mann-Whitney test, Spike patients vs. healthy (p = 0.03). (E) Box plots show the percentage of SARS-CoV-2 Spike pMHC multimer^+^ CD8^+^ T cells expressing the indicated surface markers to compare the phenotype of the HM patients and the healthy donors at TP2. Mann-Whitney test, (PD-1^+^) patients vs. healthy (p = 0.002). (F) Levels of IgG antibody against SARS-CoV-2 Spike protein subunits S1 and S2, the Spike receptor-binding domain (RBD), and nucleocapsid (N) protein in the HM patients at Pre-vac, TP1 and TP2. Threshold for antibody response was set as ≥4-fold increase in geometric mean of the MFI value for Pre-vac time point. Kruskal–Wallis one-way ANOVA with Dunn’s correction adjusting p-values with the Bonferroni method, **** (p < 0.0001), *** (p < 0.001) and ** (p < 0.01).(G) Venn diagram showing summary of the relationship between the SARS-CoV-2 Spike specific T cell responses and the IgG antibody responses in the HM patients at TP2.

## Results and discussion

### BNT162b2 mRNA vaccine induces persistent antigen-specific CD8^+^ T cells in CLL and MDS patients

BNT162b2 mRNA vaccine-induced Spike antigen-specific CD8^+^ T cells were detected in a substantial fraction of CLL and MDS patients already between day 7-10 after the first dose of vaccination (TP1). The frequency and total number of Spike-specific T cell responses further increased after the second vaccination (TP2) and persisted up to six months (TP3) following the initial vaccination (Figure 1C, Supplemental Figure 2 and Supplemental Table 8). Altogether, we identified 59 immunogenic epitopes of the 506 tested pMHC specificities across all time points. Most of the T cell responses were identified to HLA-A01:01 (10 responses), HLA-A02:01 (11 responses), and HLA-A24:02 (22 responses) restricted Spike peptides (Supplemental Figure 3A). No difference was observed in the vaccine-specific T cell responses between CLL and MDS patients (Supplemental Figure 3B). CEF epitope-specific CD8^+^ T cells were identified in samples from all four time points with no observed changes pre- to post-SARS-CoV-2 vaccination, except for a rise in influenza epitope-specific CD8^+^ T cells in few patients, likely resulting from influenza vaccination (Figure 1C, Supplemental Table 1).

Strikingly, a vaccine-driven enrichment of CD8^+^ T cells was detected in the patient group with a significantly higher frequency of Spike-specific CD8^+^ T cells compared to a healthy donor cohort evaluated at a similar time point after the second vaccination (Figure 1D, Supplemental Figure 4, and Supplemental Table 9). Also, the number of Spike-derived epitopes recognized in the patient group appeared higher, although not significant (Figure 1D). Overall, after two doses of vaccination (at TP2), Spike-specific CD8^+^ T cells were detected in 17 out of 28 patients and 10 out of 19 of healthy donors (Supplemental Figures 2, 4). Phenotypic characterization, based on the cell surface markers (CD39, CD69, HLA-DR, and PD-1; Supplemental Figure 5A) of multimer^+^ SARS-CoV-2 Spike-specific CD8^+^ T cells, displayed a signature of T cell activation in both cohorts, while a significantly higher fraction of such cells expressed PD-1 in the healthy donors (Figure 1E).

We next compared the T cell immunity with the vaccine-driven humoral immunity by analyzing plasma levels of IgG, IgM, and IgA antibodies. In contrast to the early detection of vaccine-induced T cells (Figure 1C) the antibody levels remained low at TP1, and a significant increase was observed only after the second vaccination at TP2 (Figure 1F, Supplemental Figure 6A). We found no association between the level of vaccine-specific T cells and the level of antibody response (Supplemental Figure 6B); 41% of the cohort had both T cell and antibody response, compared to 22% and 30% of individuals with only T cell response or with only antibody response, respectively (Figure 1G).

### Booster vaccination maintains vaccine induced long-term CD8^+^ T cell memory

Once established, the vaccine induced SARS-CoV-2 antigen-specific T cell immunity remained long-term (at TP3, six-months post-vaccination) with a median of one T cell response and the frequency of multimer^+^ T cells reaching up to 0.2% of the total CD8^+^ T cells in blood (Figure 2A). Importantly, a booster vaccination was required to maintain a higher frequency of vaccine-induced CD8^+^T cells. Patients without a booster vaccination (n=11) showed an overall decline in the Spike antigen-specific CD8^+^ T cell frequency from TP2 to TP3, whereas a significantly higher frequency, and breadth (total number of Spike-epitope recognized), was observed in patients vaccinated with a booster dose (n=17) prior to TP3 (Figure 2B, C), suggesting a positive impact of the booster vaccine in this patient group.

**Figure 2.**
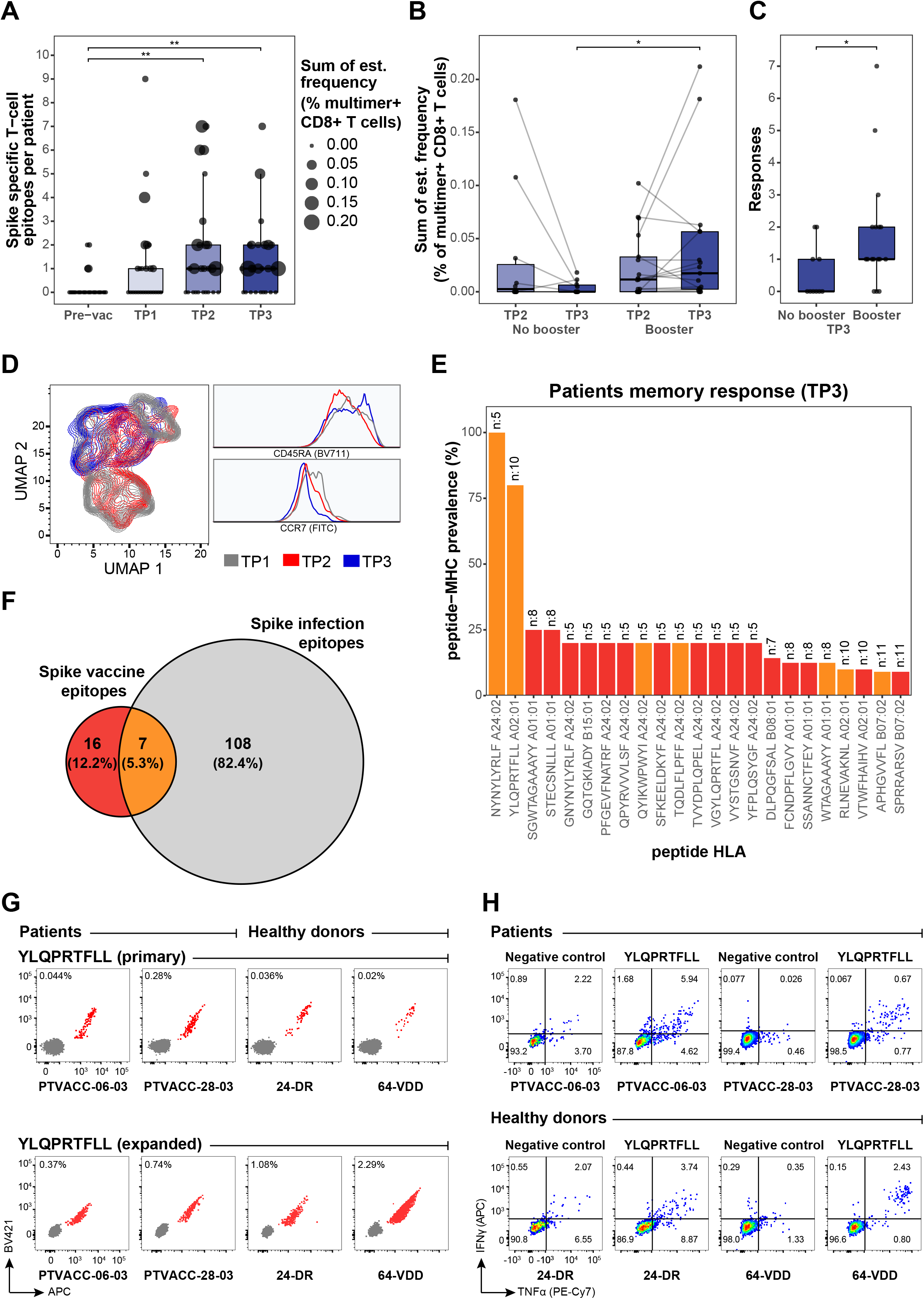
Vaccine-derived immunogenic epitopes establish functional and long-term T cell memory. (A) Summary of the number of SARS-CoV-2 Spike specific T epitopes per individual in the HM patients before (Pre-vac) and after vaccination at TP1, TP2 and TP3. The size of each dot is proportional to the sum of the estimated frequencies (%) for the significant responses in each individual. Kruskal–Wallis one-way ANOVA with Dunn’s correction adjusting p-values with the Bonferroni method, Pre-vac vs. TP2 (p = 0.001) and Pre-vac vs. TP3 (p = 0.002).(B) Box plot comparing the estimated frequencies of the SARS-CoV-2 Spike-specific CD8^+^ T cells at TP2 and TP3 between HM patients vaccinated (booster; n=17) or not vaccinated (No booster; n=11) with a booster dose before TP3. Mann-Whitney test, TP3 no booster vs. TP3 booster (p = 0.02).(C) Box plot comparing the number of SARS-CoV-2 Spike-derived CD8^+^ T cell responses between booster and non-booster HM patients at TP3. Mann-Whitney test, no booster vs. booster (p = 0.04). (D) UMAP overlay (left) of SARS-CoV-2 Spike multimer^+^ T cells showing the clustering distribution at TP1, TP2 and TP3 according to the cell’s phenotype profile. Histogram overlay (right) comparing the expression of CD45RA and CCR7 cell surface markers in the SARS-CoV-2 Spike multimer^+^ T cells at TP1, TP2 and TP3 in the HM patients. (E) Prevalence of the vaccine-induced CD8^+^ T cell memory towards SARS-CoV-2 Spike epitopes detected in the HM patients at six-months (TP3) post-vaccination. The number at the top of the bar represents the total number of patients analyzed for the corresponding pMHC specificity. Bars are colored according to the Venn diagram in the panel F.(F) Venn diagram showing the number of CD8^+^ T cell epitopes unique to the BNT162b2 mRNA vaccine (red), and SARS-CoV-2 infection (grey) and those identified in both cases (orange). Numbers were obtained from the epitopes listed in Supplemental Table 10.(G) Combinatorial tetramer analysis of HLA-A02:01 restricted YLQPRTFLL immunodominant epitope in two HM patients (at TP3) and two healthy donors (at TP2) from PBMCs, directly stained *ex vivo* (top) and after *in vitro* expansion of PBMCs incubated with YLQPRTFLL peptides for two weeks (bottom). The number in the plot shows the percentage of CD8^+^ T cells binding to pMHC tetramers.(H) Functional evaluation of the Spike immunodominant epitope YLQPRTFLL-specific CD8^+^ T cells expanded from PBMCs of two HM patients (at TP3) and two healthy donors (at TP2) by intracellular cytokine staining after peptide stimulation and no stimulation (negative). Flow cytometry plots indicates the frequency (%) of CD8^+^T cells double or single positive for TNF-α and INF-γ.

Post-vaccination longitudinal analysis of antigen-specific CD8^+^T cells phenotype in HM patients further revealed an early activation of vaccine-induced T cells (Supplemental Figure 5B) followed by a clear transition to circulating memory T cells (CD45RA^+^ CCR7^-^) (Figure 2D), which could be essential for T cell mediated long-term protection.^17,18^ At six-months postvaccination, the long-term memory was established by a total of 23 Spike-derived epitopes and a strong immunodominance was observed for NYNYLYRLF (100% prevalence) and YLQPRTFLL (80% prevalence) epitopes restricted to HLA-A24:02 and HLA-A02:01 respectively (Figure 2E, Supplemental Figure 7). These two epitopes were among the 7 immunogenic peptides that showed T cell activation after vaccination and natural infectionn,^19–23^ thus, likely to provide immune protection in SARS-CoV-2 infection (Figure 2F).^24,25^ Furthermore, even with an overall narrow CD8^+^ T cell repertoire compared to natural infection, 16 of the Spike-derived immunogenic epitopes were vaccine-unique across the nine HLAs tested in this study (Figure 2F, Supplemental Table 10) and requires further evaluation of their impact in disease protection.

To evaluate effectiveness of antigen-specific memory T cells, we validated *ex vivo* frequencies of CD8^+^ T cells reactive to several Spike and CEF epitopes using fluorophore labelled pMHC tetramers (Supplemental Figure 8) and showed comparable antigen-specific expansion of vaccine-induced CD8^+^T cells in HM patients and healthy donors (Figure 2G). The expanded YLQPRTFLL-HLA-A02:01-reactive CD8^+^ T cells were functionally active upon peptide stimulation and the cytokine secretion profiles were comparable between HM patients and healthy donors (Figure 2H, Supplemental Figure 9). Altogether, our data shows presence of functionally active long-lasting memory CD8^+^ T cells in HM patients vaccinated with the BNT162b2 mRNA vaccine.

## Acknowledgments

We thank all patients and healthy donors for participating and contributing the analyzed samples; B. Rotbøl, A. F. Løye, and A. G. Burkal for excellent technical support, and all the collaborators for active participation to this work. This work is supported by the Independent Research Fund Denmark (grant no. 0214-00053B, 2020), DFF project 1 for health and disease (grant no. 0134-00390B) to S.R.H., and by Danish Cancer Society (grant no. R306-A18139) to S.K.S.

## Authorship

S.P.A.H. designed and performed the experiments, analyzed the data, and wrote the manuscript. D.S.H, designed the research, facilitated, and collected patient samples, analyzed clinical data, and wrote the manuscript. K.K.M. analyzed the data, prepared the figures, and wrote the manuscript. T.T, and D.T. designed and performed experiments. M.T. coordinated and facilitated clinical samples. A.O.G, and L.B. supervised the clinical study, patient participation, clinical data and sample collection, and wrote the manuscript. S.R.H conceived the idea, designed, and supervised the study, and wrote the manuscript. S.K.S., conceived the idea, designed and supervised the study, designed and performed the experiments, analyzed the data, and wrote the manuscript.

## Conflicts of interest

SRH is the cofounder of PokeAcell and is the co-inventor of the patents WO2015185067 and WO2015188839 for the barcoded MHC technology which is licensed to Immudex and co-inventor of the licensed patent for Combination encoding of MHC multimers (EP2088/009356), licensee: Sanquin, NL. All other authors declare that they have no competing interests.

