## Supplementary data for "Three-doses of BNT162b2 COVID-19 mRNA vaccine establishes long-lasting CD8^+^ T cell immunity in CLL and MDS patients"

**Affiliation of Institutions**

^1^Department of Health Technology, Section of Experimental and Translational Immunology, Technical University of Denmark, Kongens Lyngby, Denmark

^2^Department of Hematology, University Hospital of Copenhagen, Rigshospitalet, Copenhagen, Denmark

^3^Innovative Immunological Models Unit, National Cancer Institute Pascale Foundation – IRCCS, Napoli, Italy

*Equal contribution

**Supplemental Methods**

**Clinical samples**

The patient cohort consist of 28 hematological patients: 23 with chronic lymphocytic leukemia (CLL) and 5 with myelodysplastic syndrome (MDS). Patients received either two doses of the BNT162b2 COVID-19 mRNA vaccine (no booster; n = 11) or two vaccine doses and a booster (booster; n = 17). The patients were screened and selected by attending physicians with the Inclusion criteria of age above 18 years, a diagnosis of CLL or MDS, individuals who were prioritized for vaccination and who did not receive ongoing treatment affecting the immune system, and with no prior COVID-19 infection. Patients’ medical record and vaccination schedule are shown in Supplemental Table 1. Approval for the study design and sample collection was obtained from the Committee on Health Research Ethics in the Capital Region of Denmark. After receiving oral and written information about the project, all patients gave their informed written consent for inclusion in the study. Blood samples were collected at four timepoints: prior to vaccination (Pre-vac), 7 to 10 days after first vaccine dose (TP1), 2 months after first vaccination dose (TP2), and 6 months after first vaccine dose (TP3). Samples from healthy donors (n=19; Supplemental Table 2) who received two doses of the BNT162b2 mRNA vaccine were collected 2 months after first vaccine dose and obtained from the National Cancer Institute Foundation Pascale, Naples, Italy. The study was approved by the institutional ethics committee with the inclusion criteria of age between 18 and 70 years.

**PBMC isolation and HLA genotyping**

Peripheral blood mononuclear cells (PBMC) were isolated from blood samples by density gradient centrifugation using Leucosep tubes (Greiner Bio-One 227288) with Lymphoprep media (StemCell Technologies 07861), and thereafter cryopreserved in Fetal Bovine Serum (FCS; Gibco 10500-064) with 10% DMSO until analysis. PBMC samples from HM patients were HLA genotyped for HLA-A, B, and C loci (IMGM Laboratories GmbH, Germany, next-generation sequencing) (Supplemental Table 5). Healthy donor samples were HLA genotyped for HLA-A loci (Laboratorio di Istoconpatibilità, UOSD Criopreservazione e BaSCO, AORN Santobono-Pausilipon, Naples, Italy) (Supplemental Table 6).

**Peptides**

All peptides used in the analysis were custom synthesized by Pepscan (Pepscan Presto BV, Lelystad, Netherlands), dissolved to 10 mM dimethyl sulfoxide (DMSO; Sigma Aldrich D2650) and stored at -20°C until use.

**MHC class I monomer production**

MHC class I monomers were produced as previously described^1,2^. Briefly, the MHC heavy chain of the selected HLA types and human ß2-microglobulin (hß2m) light chain were separately produced as inclusion bodies in *Escherichia coli* strain BL21(DE3)pLysS (Novagen 69451) using pET series expression plasmids. Solubilized heavy chain and hβ2m proteins were combined and folded *in vitro* to form functionally active HLA monomers using UV-sensitive HLA specific peptide ligands^2^. HLA-A02:01 and A24:02 molecules were folded and purified empty, as described previously^3^. Folded monomers were biotinylated with BirA biotin-protein ligase standard reaction kit (Avidity LLC, Aurora, Colorado), and purified using size exclusion-high-performance liquid chromatography (SEC-HPLC) with a BioSuite preparative chromatography column (Waters Corporation, USA) and stored in -80°C until further use.

**DNA-barcoded pMHC multimer library preparation**

DNA-barcoded multimer libraries for SARS-CoV-2 Spike peptides were generated as previously described by Bentzen et al^4^. Briefly, individual peptide–MHC (pMHC) complexes were generated by incubating for 1 hour 200 μM of each peptide with 100 μg/mL of their respective MHC molecules using direct peptide loading^3^ for HLA-A02:01 and A24:02 or by UV-mediated peptide exchange^2^ for the other HLAs. The pMHC monomers were then coupled to an allophycocyanin (APC)- and DNA barcode-labeled dextran backbone to provide each pMHC complex a unique DNA barcode label together with an APC-fluorescent label to be able to identify antigen-specific T cell populations. A DNA-barcoded pMHC multimer library for CEF peptides was generated similarly using phycoerythrin- (PE) conjugated dextran attached with unique barcodes. DNA-barcoded multimers were then used for detection of pMHC-specific T cells.

**T cell detection with DNA-barcoded pMHC multimers and phenotype panel**

PBMC from both cohorts were thawed in RPMI (Gibco 72400021) + 10% fetal bovine serum (FCS; Gibco, 10500064) + 100 µg/ml DNAse I (StemCell Technologies 07470) + 5mM MgCl_2_ and washed twice in RPMI + 10% FCS. For CLL patient samples, CD3^+^ T cells were isolated from PBMCs by immunomagnetic negative selection using the EasySep Human T-cell isolation kit as per the manufacturer’s instructions (StemCell Technologies 17951). For MDS patients and healthy cohort samples, PBMCs were used for T cell staining without isolation. Cells from all samples were then washed once in barcode-cytometry buffer (BCB; PBS + 0.5% BSA + 100 μg/mL herring DNA + 2 mM EDTA).

Patient and healthy donor PBMC samples were incubated with HLA-matching SARS-CoV-2 Spike and CEF DNA-barcoded pMHC multimers for 15 min at 37 °C, followed by incubation at 4 ˚C for 30 min with a phenotype antibody panel (Supplemental Table 7). Cells were washed twice with BCB, fixed in 1% paraformaldehyde (PFA), washed twice more, and resuspended in BCB. Cells were then acquired on a flow cytometry (AriaFusion, BD Biosciences) and pMHC multimer binding CD8^+^ T cells were sorted (Supplemental Figure 1). Sorted cells were centrifuged for 10 min at 5000 × g and the supernatant was discarded with minimal residual volume.

**DNA-barcode sequence analysis**

DNA barcodes from the isolated cells as well as from an aliquot of the multimer pool (10,000x final dilution in the PCR reaction; used as a baseline) were PCR amplified using the Taq PCR Master Mix Kit (Qiagen, 201443) and a sample-specific forward primer containing an A-key as sample identifier. Amplified barcodes were purified using the QIAquick PCR Purification kit (Qiagen 28104) and sequenced at PrimBio (USA) using an Ion Torrent PGM 316 or 318, or an Ion S5 530 chip (Life Technologies).

DNA barcode sequencing data was processed using the Barracoda software package^4^ (<https://services.healthtech.dtu.dk/service.php?Barracoda-1.8>). This tool identifies the DNA barcodes used in an experiment, assigns a sample ID and pMHC specificity to each barcode, calculates the number of reads and clonally reduced reads for each pMHC-associated DNA barcode, and includes statistical processing of the data. Fold change (FC) in read counts mapped to a given sample relative to the mean read counts mapped to triplicate baseline samples are estimated using normalization factors determined by the trimmed mean of M-values method. P-values were calculated by comparing each experiment individually to the mean baseline sample reads using a negative binomial distribution with a fixed dispersion parameter set to 0.1. False-discovery rates (FDRs) were estimated using the Benjamini–Hochberg method. At least 1/1,000 reads associated with a given DNA barcode relative to the total number of DNA barcode reads in that given sample was set as threshold to avoid false-positive detection of T cell responses. DNA barcodes with FDR < 0.1% (corresponding to p < 0.001) and Log_2_FC >2 over the baseline values for the total pMHC library were considered significant and to be true T cell responses.

The T cell frequency for each significantly enriched barcode was calculated from the percentage read count of the barcode out of the percentage of CD8^+^ multimer^+^ T cells. A non-HLA-matching and unvaccinated healthy donor was included as a negative control and any T cell recognition determined in this sample was removed from the full data set to exclude potential non-specific pMHC binding to T cells.

**T cell staining with fluorescently-labelled pMHC tetramers**

pMHC complexes with a T cell response detected using the DNA-barcode labeled multimers were selected to generate combinatorial fluorescently labeled pMHC tetramers as previously described^5,6^. Single-fluorochrome pMHC tetramers were produced by conjugating individual pMHC complexes, generated as described above, to a library of fluorophore-labeled streptavidin (SA) molecules consisting of PE-SA (Biolegend 405204), PE-CF594-SA (BD Biosciences 562284), APC-SA (Biolegend 405207) BUV395-SA (BD Biosciences 564176), BV421-SA (BD Biosciences 563259), BV737 (BD Biosciences 564293). pMHC molecules were incubated with their respective SA-conjugated fluorochrome for 30 min at 4°C, followed by incubation with D-biotin (Avidity LCC, Bio200) (25 μM final concentration) for 20 min at 4°C. pMHC tetramers for each specificity were generated in two colors and mixed in a 1:1 ratio before staining the cells.

PBMCs from HD and HM patient cohorts were expanded in vitro using a single peptide (1 µM) and cytokines IL-2 (100 IU/mL) and IL-15 (25 IU/mL) for 2 weeks in X-vivo media (Lonza BE02-060Q) with 5% human serum (HS; Gibco 1027-106). Expanded cells were used to measure peptide-specific T cell activation or stained using pMHC tetramers to detect T cells recognizing SARS-CoV-2 or CEF epitopes.

Primary and expanded PBMCs from both cohorts were thawed and washed in RPMI + 10% FCS. Cells were incubated with desatinib (50 nM final concentration) and 1 μL of pooled pMHC multimers per specificity for 15 min at 37°C in 80 μL total volume. Cells were then mixed with 20 μL antibody staining solution containing CD8-BV480 (BD Biosciences B566121) (final dilution 1/50), dump channel antibodies (CD4-FITC (BD 345768; final dilution 1/80), CD14-FITC (BD 345784; final dilution 1/32), CD19-FITC (BD Biosciences 345776; final dilution 1/16), CD40-FITC (Serotech MCA1590F; final dilution 1/40), CD16-FITC (BD Biosciences 335035; final dilution 1/64)) and a dead cell marker (LIVE/DEAD Fixable Near-IR (Invitrogen L34976; final dilution 1/1000)) and incubated for 30 min at 4°C. Cells were washed twice in FACS buffer (PBS+2% FCS) and acquired on a LSRFortessa flow cytometer (BD Biosciences).

**T cell functional analysis**

The functional capacity of T cells was measured using intracellular cytokines IFN-γ and TNF-α upon stimulation with specific peptides. Expanded PBMCs, generated as described above, from 2 healthy donors and 2 HM patients were incubated in X-vivo + 5% HS + protein transport inhibitor (GolgiPlug; BD Biosciences 555029; final dilution 1/1000) and stimulated with 1 µM of SARS-CoV-2 Spike single epitope for 8 hours at 37°C, 5% CO_2_. Cells incubated with Leukocyte Activation Cocktail (BD 550583; final dilution 1/500) were used as a positive control, and cells incubated with DMSO (final dilution 1/10,000) were used as negative control for each donor sample. Cells were stained with the surface marker antibodies CD3-FITC (BD Biosciences 345764; final dilution 1/20), CD4-BUV395 (BD Biosciences 563550 (final dilution 1/300), CD8-BV480 (BD Biosciences 566121 (final dilution 1/50)), and dead cell marker (LIVE/DEAD Fixable Near-IR (Invitrogen L34976; final dilution 1/1000)) and incubated for 30 min, 4 °C. Cells were washed twice in FACS buffer, incubated 20 min, 4 °C in fixation buffer (eBioscience 00-8222-49), washed and resuspended in permeabilization buffer (1:10 buffer to water, eBioscience 00-8333-56). Cells were stained with the intracellular antibodies PE-Cy7-TNFα (BioLegend 502930; final dilution 1/20) and APC-IFNγ antibody (BD 341117; final dilution 1/20) and incubated 30 min, 4 °C. Cells were washed twice in permeabilization buffer and resuspended in FACS buffer. CD8^+^ T cells producing intracellular cytokines were acquired on a LSRFortessa flow cytometer (BD Bioscience).

**Flow cytometry analysis**

All flow cytometry data was analyzed using FlowJo data analysis software (version 10.8.1; FlowJo LLC). For phenotype analysis of cells stained with DNA-barcoded multimers, we gated on single, live, CD8^+^, SARS-CoV-2 Spike multimer positive lymphocytes (APC^+^) and CEF multimers positive lymphocytes (PE^+^), and calculated the frequencies for specific cell populations of the multimer positive gates (Supplemental Figure 1). For UMAP analysis^7^ of SARS-CoV-2 Spike multimers-specific T cells, FCS files of samples from the patient cohorts were concatenated at the APC-positive population gate, and visualized using UMAP analysis (Version 3.1, FLowJo plugin) based on the selected markers; CD38, CD39, CD69, CD137, HLA-DR, PD-1, CCR7, CD45RA, and CD27. For antigen-specific T cell identification using combinatorial tetramer staining, we gated on single, live, CD8^+^ and FITC^-^ (dump channel) lymphocytes and selected cells positive in two tetramer colors and negative in the remaining colors (Supplemental Figure 8A)^6^. For functional evaluation, we gated on single, live, CD8^+^ lymphocytes and calculated the frequency (%) of CD8^+^ T cells double or single positive for the analyzed cytokines (Supplemental Figure 9).

**Multiplex immunoassay for antibody detection**

The multiplex bead-binding assays Milliplex SARS-CoV-2 Antigen Panels (Millipore HC19SERM1-85K, HC19SERG1-85K, HC19SERA1-85K) were used to test immunoglobulin antibody levels (IgM, IgG, and IgA) against SARS-CoV-2 Spike protein subunits S1 and S2, the Spike receptor-binding domain (RBD), and nucleocapsid (N) protein. Plasma collected during PBMC isolation was centrifuged for 10 minutes at 1000 x g and processed as per the manufacturer’s instructions. Samples were analyzed using a Bio-Plex MAGPIX Multiplex Reader (Bio-Rad Laboratories) in combination with xPotent software version 4.2 (Luminex Corporation). Antibody levels were determined as MFI (Mean Fluorescence Intensity).

**Data processing and statistical analysis**

T cell recognition data, determined by DNA-barcoded pMHC multimers analysis and Barracoda software, was plotted using RStudio version 4.1.0^8^ where peptide sequences with no significant enrichments are shown as gray dots and all peptide with a negative enrichment are set to LogFC equal zero (Figures 1C, Supplemental Figures 2 and 4). RStudio was also used to generate heatmap (Supplemental Figure 7), scatter (Supplemental Figure 6B) and box plots (Figures 1D-F and 2A-C, Supplemental Figures 3A-B, 5A and 6A) for data visualization. For the statistical analysis, data were assumed to have a non-Gaussian distribution and non-parametric tests were therefore used. For single-paired and unpaired comparisons, we used the Mann-Whitney test. For multiple comparisons, we used Kruskal–Wallis one-way ANOVA with Dunn’s correction adjusting p-values with the Bonferroni method. All statistical tests were performed using RStudio and p-values are indicated in figure legends.

**Supplemental Tables**

**Supplemental Table 1. Patient cohort information**

| **S.No.** | **Patient ID** | **Age** | **Gender** | **Diagnosis** | **Current treatment** | **Previous treatment** | **Comorbidity** | **BNT162b2 COVID-19 vaccine** | | | **Influenza vaccine** | | **Blood sample collection** | | | |
| --- | --- | --- | --- | --- | --- | --- | --- | --- | --- | --- | --- | --- | --- | --- | --- | --- |
|  |  |  |  |  |  |  |  | **Dose 1** | **Dose 2** | **Booster** | **Date** | **Type** | **Pre-vac** | **TP1** | **TP2** | **TP3** |
| 1 | PTVACC-01 | 65 | M | CLL |  |  | Previous prostate cancer, AIHA. | 26-04-21 | 25-05-21 | 11-09-21 |  |  | 04-03-21 | 21-05-21 | 25-06-21 | 04-11-21 |
| 2 | PTVACC-02 | 70 | M | CLL |  |  |  | 12-04-21 | 08-05-21 |  | 01-10-20 | Influvactetra | 04-03-21 | 22-04-21 | 11-06-21 | 12-10-21 |
| 3 | PTVACC-03 | 66 | F | CLL |  |  |  | 13-03-21 | 09-04-21 |  | 14-10-20 | Vaxigriptetra | 04-03-21 | 22-03-21 | 28-05-21 | 14-09-21 |
| 4 | PTVACC-04 | 76 | M | CLL |  |  |  | 13-04-21 | 04-05-21 | 11-09-21 | 08-10-20 | Influvactetra | 04-03-21 | 26-04-21 | 11-06-21 | 12-10-21 |
| 5 | PTVACC-05 | 75 | M | CLL | Subsitution of gammaglobulin every 10th day during the winter | Rituximab (2013) due to AIHA |  | 06-03-21 | 01-04-21 |  | 05-10-20 | Influvactetra | 04-03-21 | 15-03-21 | 10-05-21 | 14-09-21 |
| 6 | PTVACC-06 | 79 | M | CLL |  | Rectal cancer (2012), treated with RT, CT and surgery. Apoplexy (2020) |  | 19-03-21 | 09-04-21 | 15-09-21 | 26-10-20 | Vaxigriptetra | 04-03-21 |  | 07-05-21 | 06-10-21 |
| 7 | PTVACC-07 | 78 | F | CLL |  |  |  | 19-03-21 | 09-04-21 |  | 08-10-20 | Influvactetra | 04-03-21 | 26-03-21 | 21-05-21 | 14-09-21 |
| 8 | PTVACC-08 | 73 | F | CLL | Subsitution of gammaglobulin every 10th day during the winter |  |  | 11-04-21 | 05-05-21 | 11-09-21 | 04-11-20 | Influvactetra | 04-03-21 | 19-04-21 | 11-06-21 | 12-10-21 |
| 9 | PTVACC-09 | 76 | F | CLL |  |  |  | 13-04-21 | 08-05-21 | 15-09-21 | 07-10-20 | Influvactetra | 04-03-21 | 22-04-21 | 18-06-21 | 12-10-21 |
| 10 | PTVACC-10 | 83 | M | CLL |  |  |  | 21-03-21 | 14-04-21 |  | 07-10-20 | Vaxigriptetra | 04-03-21 |  |  | 24-09-21 |
| 11 | PTVACC-12 | 68 | F | CLL |  |  |  | 05-03-21 | 31-03-21 |  |  |  | 04-03-21 | 15-03-21 | 07-05-21 | 14-09-21 |
| 12 | PTVACC-13 | 83 | F | CLL |  |  | Diabetes, previous breast cancer. | 07-03-21 | 02-04-21 |  |  |  | 04-03-21 | 15-03-21 | 10-05-21 | 14-09-21 |
| 13 | PTVACC-14 | 62 | M | CLL |  |  |  | 12-05-21 | 23-06-21 | 03-11-21 | 05-10-21 | Vaxigriptetra | 10-03-21 | 19-05-21 | 13-07-21 | 16-11-21 |
| 14 | PTVACC-16 | 73 | M | MDS (CMML) | Deferasirox. Regular blood transfusions. |  |  | 10-04-21 | 05-05-21 | 13-09-21 | 09-10-20 | Vaxigriptetra | 10-03-21 | 19-04-21 | 18-06-21 | 07-10-21 |
| 15 | PTVACC-17 | 74 | F | MDS | Deferasirox. Regular blood transfusions. |  |  | 18-04-21 | 13-05-21 | 11-09-21 | 05-10-20 | Influvactetra | 10-03-21 | 29-04-21 | 18-06-21 | 26-10-21 |
| 16 | PTVACC-18 | 83 | M | CLL |  |  |  | 18-03-21 | 12-04-21 |  | 09-10-20 | Vaxigriptetra | 10-03-21 | 26-03-21 | 04-06-21 | 24-09-21 |
| 17 | PTVACC-19 | 69 | F | MDS | Deferasirox. Regular blood transfusions. |  | Azacitidine, erythropoietin. | 10-04-21 | 01-05-21 | 13-09-21 |  |  | 10-03-21 | 19-04-21 | 18-06-21 | 07-10-21 |
| 18 | PTVACC-20 | 73 | F | MDS |  | Interferon |  | 18-04-21 | 13-05-21 |  | 09-10-20 | Vaxigriptetra | 10-03-21 | 26-04-21 | 25-06-21 | 16-11-21 |
| 19 | PTVACC-21 | 78 | F | MDS | Deferasirox. Regular blood transfusions. | Revlimid |  | 08-04-21 | 06-05-21 |  | 02-10-20 | Influvactetra | 10-03-21 | 21-04-21 | 16-06-21 | 06-10-21 |
| 20 | PTVACC-23 | 78 | M | CLL |  | Rituximab + Bendamustin (last treatment sep 2020) | Diabetes | 29-03-21 | 20-04-21 | 12-09-21 |  |  | 17-03-21 | 07-04-21 | 28-05-21 | 24-09-21 |
| 21 | PTVACC-25 | 70 | F | CLL |  |  |  | 18-04-21 | 09-05-21 | 10-09-21 | 07-10-20 | Vaxigriptetra | 22-03-21 | 26-04-21 | 11-06-21 | 07-10-21 |
| 22 | PTVACC-26 | 54 | M | CLL |  |  |  | 25-05-21 | 01-07-21 | 17-09-21 |  |  | 09-04-21 | 04-06-21 | 12-08-21 | 16-11-21 |
| 23 | PTVACC-28 | 67 | M | CLL |  |  |  | 27-04-21 | 01-06-21 | 22-09-21 | 16-10-20 | Vaxigriptetra | 16-04-21 | 04-05-21 | 25-06-21 | 02-11-21 |
| 24 | PTVACC-29 | 59 | M | CLL |  |  |  | 11-05-21 | 16-06-21 | 03-11-21 |  |  | 23-04-21 | 19-05-21 | 13-07-21 | 16-11-21 |
| 25 | PTVACC-30 | 74 | M | CLL |  |  |  | 10-04-21 | 04-05-21 | 14-09-21 | 12-10-20 | Influvactetra | 09-04-21 | 19-04-21 | 08-06-21 | 07-10-21 |
| 26 | PTVACC-31 | 62 | F | CLL |  |  |  | 09-05-21 | 16-06-21 | 23-09-21 |  |  | 16-04-21 | 19-05-21 | 09-07-21 | 16-11-21 |
| 27 | PTVACC-32 | 72 | F | CLL |  | Rituximab + Bendamustin (2019) |  | 17-04-21 | 09-05-21 | 14-09-21 | 02-10-20 | Vaxigriptetra | 14-04-21 | 26-04-21 | 21-06-21 | 07-10-21 |
| 28 | PTVACC-33 | 71 | F | CLL |  |  |  | 17-04-21 | 08-05-21 |  | 07-10-20 | Influvactetra | 16-04-21 | 26-04-21 | 18-06-21 | 12-10-21 |

**Supplemental Table 2. Healthy donors information**

| **S. No.** | **Healthy donor ID** | **Gender** | **BNT162b2 COVID-19 vaccine** | | **Blood sample collection** |
| --- | --- | --- | --- | --- | --- |
|  |  |  | **Dose 1** | **Dose 2** | **TP2** |
| 1 | 3-GS | M | 27-12-20 | 17-01-21 | 28-01-21 |
| 2 | 4-FM | M | 27-12-20 | 17-01-21 | 02-02-21 |
| 3 | 7-RDF | F | 31-12-20 | 21-01-21 | 05-02-21 |
| 4 | 9-PM | M | 31-12-20 |  | 20-01-21 |
| 5 | 10-PA | F | 01-01-21 | 22-01-21 | 08-02-21 |
| 6 | 12-BV | F | 02-01-21 | 23-01-21 | 10-02-21 |
| 7 | 13-GB | F | 05-01-21 | 26-01-21 | 12-02-21 |
| 8 | 15-CR | F | 08-01-21 | 29-01-21 | 10-02-21 |
| 9 | 21-AI | M | 08-01-21 | 28-01-21 | 08-02-21 |
| 10 | 24-DR | F | 08-01-21 | 29-01-21 | 15-02-21 |
| 11 | 25-CE | F | 08-01-21 | 29-01-21 | 12-02-21 |
| 12 | 27-ACP | F | 10-01-21 | 30-01-21 | 15-02-21 |
| 13 | 32-MT | F | 13-01-21 |  | 15-02-21 |
| 14 | 36-DDA | F | 11-01-21 |  | 29-01-21 |
| 15 | 47-SA | M | 13-01-21 | 03-02-21 | 19-02-21 |
| 16 | 48-PF | F | 13-01-21 | 03-02-21 | 19-02-21 |
| 17 | 50-AC | M | 13-01-21 | 03-02-21 | 16-02-21 |
| 18 | 59-GA | M |  | 05-02-21 | 19-02-21 |
| 19 | 64-VDD | F | 03-01-21 | 24-01-21 | 08-02-21 |

**Supplemental Table 3. SARS-CoV-2 Spike peptide library with their HLA rank score**

Supplemental Table 3 can be found in the Supplemental excel file.

**Supplemental Table 4. CEF peptide library**

| **Number** | **HLA** | **Virus** | **Peptide sequence** |
| --- | --- | --- | --- |
| 1 | HLA-A01:01 | CMV | YSEHPTFTSQY |
| 2 | HLA-A01:01 | CMV | VTEHDTLLY |
| 3 | HLA-A01:01 | Influenza | VSDGGPNLY |
| 4 | HLA-A01:01 | Influenza | CTELKLSDY |
| 5 | HLA-A01:01 | Influenza | TFEFTSFFY |
| 6 | HLA-A01:01 | Influenza | AEKPKFLPDLY |
| 7 | HLA-A02:01 | Influenza | FMYSDFHFI |
| 8 | HLA-A02:01 | Influenza | AIMDKNIML |
| 9 | HLA-A02:01 | Influenza | AIMDKNIIL |
| 10 | HLA-A02:01 | Influenza | ILGFVFLTV |
| 11 | HLA-A02:01 | Influenza | MMMGMFNML |
| 12 | HLA-A02:01 | Influenza | FNMLSTVLGV |
| 13 | HLA-A03:01 | Influenza | SIIPSGPLK |
| 14 | HLA-A03:01 | Influenza | RMVLASTTAK |
| 15 | HLA-A03:01 | Influenza | RVLSFIKGTK |
| 16 | HLA-A03:01 | Influenza | RLEDVFAGK |
| 17 | HLA-A03:01 | Influenza | KSMREEYRK |
| 18 | HLA-A11:01 | EBV | AVFDRKSDAK |
| 19 | HLA-A11:01 | CMV | GPISGHVLK |
| 20 | HLA-A11:01 | Influenza | SIIPSGPLK |
| 21 | HLA-A11:01 | Influenza | RMVLASTTAK |
| 22 | HLA-A11:01 | Influenza | RVLSFIKGTK |
| 23 | HLA-A11:01 | Influenza | RLEDVFAGK |
| 24 | HLA-A11:01 | Influenza | KSMREEYRK |
| 25 | HLA-A24:02 | EBV | RYSIFFDY |
| 26 | HLA-A24:02 | EBV | TYGPVFMCL |
| 27 | HLA-A24:02 | EBV | DYCNVLNKEF |
| 28 | HLA-A24:02 | CMV | AYAQKIFKIL |
| 29 | HLA-A24:02 | Influenza | FMYSDFHFI |
| 30 | HLA-A24:02 | Influenza | SWPDGAELPF |
| 31 | HLA-A24:02 | Influenza | ITFMQALQLL |
| 32 | HLA-A24:02 | Influenza | VETPIRNEW |
| 33 | HLA-B07:02 | CMV | TPRVTGGGAM |
| 34 | HLA-B07:02 | CMV | RPHERNGFTV |
| 35 | HLA-B07:02 | EBV | RPPIFIRLL |
| 36 | HLA-B07:02 | Influenza | AIMDKNIML |
| 37 | HLA-B07:02 | Influenza | AIMDKNIIL |
| 38 | HLA-B07:02 | Influenza | SWPDGAELPF |
| 39 | HLA-B07:02 | Influenza | TTYQRTRAL |
| 40 | HLA-B07:02 | Influenza | LPFDKPTIM |
| 41 | HLA-B07:02 | Influenza | LPFDKTTVM |
| 42 | HLA-B07:02 | Influenza | LPFEKSTVM |
| 43 | HLA-B07:02 | Influenza | LPFDKSTIM |
| 44 | HLA-B07:02 | Influenza | LPFERSTIM |
| 45 | HLA-B07:02 | Influenza | LPFERATIM |
| 46 | HLA-B07:02 | Influenza | QPEWFRNVL |
| 47 | HLA-B07:02 | Influenza | SPIVPSFDM |
| 48 | HLA-B08:01 | Influenza | ELRSRYWAI |
| 49 | HLA-B08:01 | EBV | RAKFKQLL |
| 50 | HLA-B08:01 | CMV | ELRRKMMYM |
| 51 | HLA-B08:01 | EBV | QAKWRLQTL |
| 52 | HLA-B08:01 | EBV | FLRGRAYGL |
| 53 | HLA-B08:01 | Influenza | AIMDKNIML |
| 54 | HLA-B08:01 | Influenza | AIMDKNIIL |
| 55 | HLA-B08:01 | Influenza | MMMGMFNML |
| 56 | HLA-B08:01 | Influenza | TTYQRTRAL |
| 57 | HLA-B08:01 | Influenza | LPFDKPTIM |
| 58 | HLA-B08:01 | Influenza | LPFDKTTVM |
| 59 | HLA-B08:01 | Influenza | LPFEKSTVM |
| 60 | HLA-B08:01 | Influenza | LPFDKSTIM |
| 61 | HLA-B08:01 | Influenza | LPFERSTIM |
| 62 | HLA-B08:01 | Influenza | LPFERATIM |
| 63 | HLA-B15:01 | EBV | QNGALAINTF |
| 64 | HLA-B15:01 | EBV | LEKARGSTY |
| 65 | HLA-B15:01 | Influenza | TQIQTRRSF |
| 66 | HLA-B15:01 | Influenza | KMARLGKGY |
| 67 | HLA-B15:01 | EBV | GQGGSPTAM |

**Supplemental Table 5. HLA genotype data for patient donors**

| **S. No.** | **Patient ID** | **HLA-A** | | **HLA-B** | | **HLA-C** | |
| --- | --- | --- | --- | --- | --- | --- | --- |
| 1 | PTVACC-01 | 02:01:01:01 | 24:02:01:05 | 15:01:01:01 | 18:01:01 | 03:04:01 | 07:01:01 |
| 2 | PTVACC-02 | 01:01:01:01 | 32:01:01:01 | 08:01:01 | 44:02:01 | 06:02:01 | 07:01:01:06 |
| 3 | PTVACC-03 | 01:01:01:01 | 01:01:01:01 | 07:02:01 | 08:09 | 07:01:01 | 07:02:01 |
| 4 | PTVACC-04 | 03:01:01:01 | 25:01:01:01 | 07:02:01 | 18:01:01 | 07:02:01 | 12:03:01 |
| 5 | PTVACC-05 | 03:01:01:01 | 25:01:01:01 | 07:02:01 | 18:01:01 | 07:02:01 | 12:03:01 |
| 6 | PTVACC-06 | 02:01:01:01 | 11:01:01:01 | 07:02:01 | 39:24:01 | 07:01:01 | 07:02:01 |
| 7 | PTVACC-07 | 01:01:01:01 | 03:01:01:01 | 07:02:01 | 35:03:01 | 04:01:01 | 07:10 |
| 8 | PTVACC-08 | 24:02:01:01 | 24:02:01:01 | 07:02:01 | 40:01:02 | 03:04:01 | 07:02:01 |
| 9 | PTVACC-09 | 02:01:01:01 | 02:01:01:01 | 13:02:01 | 27:05:02 | 01:02:01 | 06:02:01 |
| 10 | PTVACC-10 | 01:01:01:01 | 33:01:01:01 | 08:01:01 | 14:02:01:01 | 07:01:01 | 08:02:01 |
| 12 | PTVACC-12 | 02:01:01:01 | 03:01:01:01 | 15:01:01:01 | 49:01:01 | 03:04:01 | 07:01:01 |
| 13 | PTVACC-13 | 01:01:01:01 | 31:01:02:01 | 08:01:01 | 40:01:02 | 03:04:01 | 07:01:01 |
| 14 | PTVACC-14 | 01:01:01:01 | 31:01:02:01 | 08:01:01 | 37:01:01:01 | 06:02:01 | 07:01:01 |
| 16 | PTVACC-16 | 01:01:01:01 | 11:01:01:01 | 08:01:01 | 27:05:02 | 02:02:02 | 07:01:01 |
| 17 | PTVACC-17 | 26:01:01:01 | 31:01:02:01 | 07:02:01 | 14:01:01:01 | 07:02:01 | 08:02:01:02 |
| 18 | PTVACC-18 | 26:01:01:01 | 29:02:01:01 | 07:02:01 | 44:03:01 | 07:02:01 | 16:01:01:01 |
| 19 | PTVACC-19 | 03:01:01:01 | 03:01:01:01 | 07:02:01 | 44:02:01 | 05:01:01 | 07:02:01 |
| 20 | PTVACC-20 | 01:01:01:01 | 03:01:01:01 | 08:01:01 | 35:01:01 | 04:01:01 | 07:01:01 |
| 21 | PTVACC-21 | 02:01:01:01 | 24:03:01:01 | 15:01:01:01 | 38:01:01 | 03:04:01 | 12:03:01 |
| 23 | PTVACC-23 | 24:02:01:01 | 26:01:01:01 | 08:01:01 | 39:06:02:01 | 07:01:01 | 07:02:01 |
| 25 | PTVACC-25 | 24:02:01:01 | 29:02:01:01 | 15:01:01:01 | 44:03:01 | 03:03:01 | 16:01:01:01 |
| 26 | PTVACC-26 | 02:01:01:01 | 02:01:01:01 | 15:01:01:01 | 40:01:02 | 03:03:01 | 03:04:01 |
| 28 | PTVACC-28 | 02:01:01:01 | 02:05:01:01 | 50:01:01:06 | 51:01:01 | 06:02:01:02 | 15:02:01 |
| 29 | PTVACC-29 | 03:01:01:01 | 30:02:01:01 | 07:02:01 | 18:01:01 | 05:01:01 | 07:02:01 |
| 30 | PTVACC-30 | 02:01:01:01 | 31:01:02:01 | 40:01:02 | 44:02:01 | 03:04:01 | 07:04:01 |
| 31 | PTVACC-31 | 02:01:01:01 | 25:01:01:01 | 39:01:01 | 50:01:01:01 | 06:02:01:02 | 07:02:01 |
| 32 | PTVACC-32 | 01:01:01:01 | 24:02:01:01 | 07:02:01 | 08:01:01 | 07:01:01 | 07:02:01 |
| 33 | PTVACC-33 | 02:01:01:01 | 02:01:01:01 | 37:01:01:01 | 57:01:01 | 06:02:01 | 06:02:01 |

**Supplemental Table 6. HLA genotype data for healthy donors**

| **S. No.** | **Healthy donor ID** | **HLA-A** | |
| --- | --- | --- | --- |
| 1 | 3-GS | 01:01 | 30:01 |
| 2 | 4-FM | 02:01 | 24:02 |
| 3 | 7-RDF | 02:01 | 24:02 |
| 4 | 9-PM | 01:01 | 03:01 |
| 5 | 10-PA | 01:01 | 23:01P |
| 6 | 12-BV | 03:01 | 24:02 |
| 7 | 13-GB | 01:01 | 32:01 |
| 8 | 15-CR | 02:01 | 11:01 |
| 9 | 21-AI | 01:01 | 02:01 |
| 10 | 24-DR | 02:01 | 11:01 |
| 11 | 25-CE | 02:01 | 11:01 |
| 12 | 27-ACP | 01:01 | 69:01 |
| 13 | 32-MT | 03:01 | 24:02 |
| 14 | 36-DDA | 03:01 | 68:01 |
| 15 | 47-SA | 02:01 | 24:02 |
| 16 | 48-PF | 01:01 | 02:01 |
| 17 | 50-AC | 03:01 | 68:02 |
| 18 | 59-GA | 01:01 | 24:02 |
| 19 | 64-VDD | 02:01 | 03:01 |

**Supplemental Table 7. Phenotype antibody panel**

| **Antibody** | **Conjugate** | **Clone** | **Dilution** | **Provider** | **Catalogue ID** |
| --- | --- | --- | --- | --- | --- |
| CD3 | BV786 | SK7 | 1/20 | BD Biosciences | 563800 |
| CD4 | BV650 | SK3 | 1/40 | BD Biosciences | 563875 |
| CD8 | BV480 | RPA-T8 | 1/50 | BD Biosciences | 566121 |
| CD45RA | BV711 | HI100 | 1/40 | BD Biosciences | 563733 |
| CCR7 | FITC | G043H7 | 1/20 | Biolegend | 353216 |
| CD27 | BV605 | O323 | 1/40 | Biolegend | 302830 |
| CD38 | BUV737 | HB7 | 1/160 | BD Biosciences | 612824 |
| CD39 | PE-CF594 | Tu66 | 1/40 | BD Biosciences | 563678 |
| CD69 | BUV395 | FN50 | 1/20 | BD Biosciences | 564364 |
| CD137 | PE-Cy5 | 4B4-1 | 1/40 | BD Biosciences | 551137 |
| HLA-DR | APC-R700 | G46-6 | 1/160 | BD Biosciences | 565127 |
| PD1 | BV421 | EH12.1 | 1/33 | BioLegend | 562516 |
| Live-Dead marker | APC-Cy7 | - | 1/1000 | Thermo Fishcer Scientific | L34976 |

**Supplemental Table 8: List of identified SARS-CoV-2 Spike and CEF-derived epitopes in each HM patient pre- and post-vaccination**

Supplemental Table 8 can be found in the Supplemental excel file.

**Supplemental Table 9: List of identified SARS-CoV-2 Spike and CEF-derived epitopes in each healthy donor**

| **Time point** | **Sample** | **SARS-CoV-2** | **CEF** | **HLA** | **Peptide** | **Est. frequency** | **Log fold**  **change** | **p** |
| --- | --- | --- | --- | --- | --- | --- | --- | --- |
| **TP2** | 4-FM | Spike |  | B07:02 | APHGVVFLHV | 0.00438965 | 2.774271135 | 4.21449E-07 |
|  | 7-RDF | Spike |  | A24:02 | NYNYLYRLF | 0.00073313 | 2.62053158 | 7.96152E-06 |
|  |  |  | EBV | B08:01 | RAKFKQLL | 1.302277433 | 2.220737918 | 0.000347742 |
|  | 10-PA | Spike |  | A01:01 | CNDPFLGVYY | 0.000294435 | 2.476331911 | 4.77679E-05 |
|  |  |  | CMV | A01:01 | VTEHDTLLY | 0.066164466 | 3.453303108 | 5.40338E-10 |
|  |  |  | CMV | A01:01 | YSEHPTFTSQY | 0.041188475 | 3.083039593 | 8.35565E-08 |
|  | 12-BV | Spike |  | A24:02 | NYNYLYRLF | 0.000279518 | 2.658187479 | 6.50962E-06 |
|  |  | Spike |  | B07:02 | KPSKRSFIEDL | 0.00079386 | 2.280902663 | 0.000120398 |
|  | 13-GB |  | CMV | A01:01 | VTEHDTLLY | 0.472221287 | 5.224792289 | 5.50166E-24 |
|  |  |  | CMV | B07:02 | TPRVTGGGAM | 1.402605853 | 6.188242637 | 5.68816E-32 |
|  |  |  | EBV | B08:01 | RAKFKQLL | 0.14739609 | 3.617217243 | 3.95271E-12 |
|  | 15-CR | Spike |  | B08:01 | VFQTRAGCL | 0.001879509 | 4.604412646 | 3.63587E-19 |
|  |  |  | EBV | A11:01 | AVFDRKSDAK | 0.015641379 | 2.40040148 | 9.63192E-05 |
|  | 21-AI |  | CMV | A01:01 | VTEHDTLLY | 0.046867729 | 2.855837105 | 4.51807E-07 |
|  | 24-DR | Spike |  | A02:01 | YLQPRTFLL | 0.003518624 | 2.661312193 | 1.7102E-06 |
|  |  |  | EBV | A11:01 | AVFDRKSDAK | 0.100966921 | 2.690882902 | 7.73563E-06 |
|  | 25-CE | Spike |  | A02:01 | YLQPRTFLL | 0.003356941 | 3.557378751 | 8.99132E-12 |
|  |  | Spike |  | B07:02 | APGQTGKIA | 0.018466541 | 5.886907856 | 1.6134E-29 |
|  | 27-ACP |  | CMV | A01:01 | VTEHDTLLY | 0.933285519 | 7.422775844 | 2.0529E-42 |
|  | 32-MT | Spike |  | A03:01 | KCYGVSPTK | 0.001940778 | 3.047260776 | 4.49303E-08 |
|  |  | Spike |  | A24:02 | NYNYLYRLF | 0.002133651 | 2.583660931 | 1.17714E-05 |
|  |  |  | EBV | B08:01 | RAKFKQLL | 0.015261876 | 4.259625859 | 3.6431E-16 |
|  |  |  | EBV | B08:01 | FLRGRAYGL | 0.010499391 | 2.281967002 | 8.7806E-05 |
|  | 50-AC | Spike |  | B07:02 | VVNQNAQAL | 0.002737023 | 2.726455031 | 8.86024E-07 |
|  |  | Spike |  | B35:01 | HADQLTPTW | 0.002822398 | 2.259543364 | 0.000273509 |
|  | 59-GA |  | Influenza | B08:01 | TTYQRTRAL | 0.049391289 | 2.223693611 | 0.000507732 |
|  | 64-VDD | Spike |  | A02:01 | YLQPRTFLL | 0.00284029 | 2.245876745 | 0.000176079 |

**Supplemental Table 10. SARS-CoV-2 Spike T cell epitopes previously reported from natural infection**.

| **HLA** | **Peptide** | **Reference** |
| --- | --- | --- |
| **A01:01** | LLTDEMIAQY | [Saini et al. 2021](https://www.science.org/doi/10.1126/sciimmunol.abf7550) |
|  | LTDEMIAQY | [Schulien et al. 2021](https://www.nature.com/articles/s41591-020-01143-2); [Nelde et al. 2021](https://www.nature.com/articles/s41590-020-00808-x); [Kared et al. 2021](https://pubmed.ncbi.nlm.nih.gov/33427749/) |
|  | **WTAGAAAYY*** | [Wagner et al. 2022](https://www.cell.com/cell-reports/fulltext/S2211-1247(21)01718-6) |
|  | YTNSFTRGVY | [Tarke et al. 2021](https://www.sciencedirect.com/science/article/pii/S266637912100015X) |
|  | YTNSFTRGVY | [Tarke et al. 2021](https://www.sciencedirect.com/science/article/pii/S266637912100015X) |
| **A02:01** | ALNTLVKQL | [Shomuradova et al. 2020](https://www.cell.com/immunity/fulltext/S1074-7613(20)30469-6) |
|  | FIAGLIAIV | [Poran et al. 2020](https://genomemedicine.biomedcentral.com/articles/10.1186/s13073-020-00767-w); [Rha et al. 2021](https://www.cell.com/immunity/fulltext/S1074-7613(20)30509-4); [Shomuradova et al. 2020](https://www.cell.com/immunity/fulltext/S1074-7613(20)30469-6); [Saini et al. 2021](https://www.science.org/doi/10.1126/sciimmunol.abf7550) |
|  | FLPFFSNV | [Saini et al. 2021](https://www.science.org/doi/10.1126/sciimmunol.abf7550?url_ver=Z39.88-2003&rfr_id=ori:rid:crossref.org&rfr_dat=cr_pub%20%200pubmed) |
|  | GLTVLPPLL | [Tarke et al. 2021](https://www.sciencedirect.com/science/article/pii/S266637912100015X); [Poran et al. 2020](https://genomemedicine.biomedcentral.com/articles/10.1186/s13073-020-00767-w) |
|  | HLMSFPQSA | [Tarke et al. 2021](https://www.sciencedirect.com/science/article/pii/S266637912100015X) |
|  | KIADYNYKL | [Chen et al. 2020](https://www.ncbi.nlm.nih.gov/pmc/articles/PMC7812294/); [Shomuradova et al. 2020](https://www.cell.com/immunity/fulltext/S1074-7613(20)30469-6) |
|  | KIADYNYKL | [Shomuradova et al. 2020](https://www.cell.com/immunity/fulltext/S1074-7613(20)30469-6) |
|  | KLNDLCFTNV | [Poran et al. 2020](https://genomemedicine.biomedcentral.com/articles/10.1186/s13073-020-00767-w) |
|  | KLPDDFTGCV | [Shomuradova et al. 2020](https://www.cell.com/immunity/fulltext/S1074-7613(20)30469-6) |
|  | KLPDDFTGCV | [Shomuradova et al. 2020](https://www.cell.com/immunity/fulltext/S1074-7613(20)30469-6) |
|  | LITGRLQSL | [Shomuradova et al. 2020](https://www.cell.com/immunity/fulltext/S1074-7613(20)30469-6) |
|  | LLFNKVTLA | [Shomuradova et al. 2020](https://www.cell.com/immunity/fulltext/S1074-7613(20)30469-6) |
|  | **RLNEVAKNL*** | [Shomuradova et al. 2020](https://www.cell.com/immunity/fulltext/S1074-7613(20)30469-6) |
|  | RLQSLQTYV | [Tarke et al. 2021](https://www.sciencedirect.com/science/article/pii/S266637912100015X); [Shomuradova et al. 2020](https://www.cell.com/immunity/fulltext/S1074-7613(20)30469-6); [Poran et al. 2020](https://genomemedicine.biomedcentral.com/articles/10.1186/s13073-020-00767-w) |
|  | SIIAYTMSL | [Tarke et al. 2021](https://www.sciencedirect.com/science/article/pii/S266637912100015X) |
|  | TLDSKTQSL | [Tarke et al. 2021](https://www.sciencedirect.com/science/article/pii/S266637912100015X); [Sekine et al. 2020](https://www.cell.com/cell/fulltext/S0092-8674(20)31008-4) |
|  | VLNDILSRL | [Habel et al. 2020](https://www.pnas.org/doi/10.1073/pnas.2015486117); [Shomuradova et al. 2020](https://www.cell.com/immunity/fulltext/S1074-7613(20)30469-6); [Saini et al. 2021](https://www.science.org/doi/10.1126/sciimmunol.abf7550) |
|  | VVFLHVTYV | [Kared et al. 2021](https://pubmed.ncbi.nlm.nih.gov/33427749/); [Tarke et al. 2021](https://www.sciencedirect.com/science/article/pii/S266637912100015X); [Saini et al. 2021](https://www.science.org/doi/10.1126/sciimmunol.abf7550) |
|  | **YLQPRTFLL*** | [Ferretti et al. 2020](https://www.ncbi.nlm.nih.gov/pmc/articles/PMC7574860/); [Tarke et al. 2021](https://www.sciencedirect.com/science/article/pii/S266637912100015X); [Shomuradova et al. 2020](https://www.cell.com/immunity/fulltext/S1074-7613(20)30469-6); [Rha et al. 2021](https://www.cell.com/immunity/fulltext/S1074-7613(20)30509-4); [Sekine et al. 2020](https://www.cell.com/cell/fulltext/S0092-8674(20)31008-4); [Habel et al. 2020](https://www.pnas.org/doi/10.1073/pnas.2015486117); [Kared et al. 2021](https://pubmed.ncbi.nlm.nih.gov/33427749/) |
| **A03:01** | ALDPLSETK | [Tarke et al. 2021](https://www.sciencedirect.com/science/article/pii/S266637912100015X) |
|  | EILPVSMTK | [Tarke et al. 2021](https://www.sciencedirect.com/science/article/pii/S266637912100015X) |
|  | GVYFASTEK | [Kared et al. 2021](https://pubmed.ncbi.nlm.nih.gov/33427749/) |
|  | GVYYHKNNK | [Tarke et al. 2021](https://www.sciencedirect.com/science/article/pii/S266637912100015X) |
|  | GVYYPDKVFR | [Tarke et al. 2021](https://www.sciencedirect.com/science/article/pii/S266637912100015X) |
|  | KCYGVSPTK | [Ferretti et al. 2020](https://www.ncbi.nlm.nih.gov/pmc/articles/PMC7574860/); [Tarke et al. 2021](https://www.sciencedirect.com/science/article/pii/S266637912100015X); [Saini et al. 2021](https://www.science.org/doi/10.1126/sciimmunol.abf7550) |
|  | KVFRSSVLH | [Tarke et al. 2021](https://www.sciencedirect.com/science/article/pii/S266637912100015X) |
|  | RASANLAATK | [Tarke et al. 2021](https://www.sciencedirect.com/science/article/pii/S266637912100015X) |
|  | RLFRKSNLK | [Tarke et al. 2021](https://www.sciencedirect.com/science/article/pii/S266637912100015X) |
|  | SVYAWNRKR | [Tarke et al. 2021](https://www.sciencedirect.com/science/article/pii/S266637912100015X) |
|  | TLADAGFIK | [Tarke et al. 2021](https://www.sciencedirect.com/science/article/pii/S266637912100015X) |
|  | TVYDPLQPELDSFK | [Tarke et al. 2021](https://www.sciencedirect.com/science/article/pii/S266637912100015X) |
|  | VTYVPAQEK | [Tarke et al. 2021](https://www.sciencedirect.com/science/article/pii/S266637912100015X) |
| **A11:01** | GTHWFVTQR | [Kared et al. 2021](https://pubmed.ncbi.nlm.nih.gov/33427749/) |
|  | GVYFASTEK | [Kared et al. 2021](https://pubmed.ncbi.nlm.nih.gov/33427749/) |
|  | RLFRKSNLK | [Kared et al. 2021](https://pubmed.ncbi.nlm.nih.gov/33427749/) |
| **A24:02** | AYSNNSIAI | [Tarke et al. 2021](https://www.sciencedirect.com/science/article/pii/S266637912100015X) |
|  | EYVSQPFLM | [Tarke et al. 2021](https://www.sciencedirect.com/science/article/pii/S266637912100015X) |
|  | GYLQPRTFLL | [Tarke et al. 2021](https://www.sciencedirect.com/science/article/pii/S266637912100015X) |
|  | HWFVTQRNF | [Tarke et al. 2021](https://www.sciencedirect.com/science/article/pii/S266637912100015X) |
|  | IYQTSNFRV | [Tarke et al. 2021](https://www.sciencedirect.com/science/article/pii/S266637912100015X) |
|  | **NYNYLYRLF*** | [Tarke et al. 2021](https://www.sciencedirect.com/science/article/pii/S266637912100015X); [Kared et al. 2021](https://pubmed.ncbi.nlm.nih.gov/33427749/) |
|  | **QYIKWPWYI*** | [Ferretti et al. 2020](https://www.ncbi.nlm.nih.gov/pmc/articles/PMC7574860/); [Tarke et al. 2021](https://www.sciencedirect.com/science/article/pii/S266637912100015X); [Nelde et al. 2021](https://www.nature.com/articles/s41590-020-00808-x); [Kared et al. 2021](https://pubmed.ncbi.nlm.nih.gov/33427749/) |
|  | RFDNPVLPF | [Tarke et al. 2021](https://www.sciencedirect.com/science/article/pii/S266637912100015X); [Kared et al. 2021](https://pubmed.ncbi.nlm.nih.gov/33427749/) |
|  | RFPNITNLCPF | [Tarke et al. 2021](https://www.sciencedirect.com/science/article/pii/S266637912100015X) |
|  | RVYSTGSNVF | [Tarke et al. 2021](https://www.sciencedirect.com/science/article/pii/S266637912100015X) |
|  | RVYSTGSNVF | [Tarke et al. 2021](https://www.sciencedirect.com/science/article/pii/S266637912100015X) |
|  | SFPQSAPHGVVF | [Tarke et al. 2021](https://www.sciencedirect.com/science/article/pii/S266637912100015X) |
|  | **TQDLFLPFF*** | [Saini et al. 2021](https://www.science.org/doi/10.1126/sciimmunol.abf7550) |
|  | TYVPAQEKNFT | [Saini et al. 2021](https://www.science.org/doi/10.1126/sciimmunol.abf7550) |
|  | VFKNIDGYF | [Tarke et al. 2021](https://www.sciencedirect.com/science/article/pii/S266637912100015X) |
|  | VFVSNGTHWF | [Tarke et al. 2021](https://www.sciencedirect.com/science/article/pii/S266637912100015X) |
|  | VYDPLQPELDSF | [Tarke et al. 2021](https://www.sciencedirect.com/science/article/pii/S266637912100015X) |
|  | VYSSANNCTF | [Tarke et al. 2021](https://www.sciencedirect.com/science/article/pii/S266637912100015X) |
|  | VYYPDKVF | [Tarke et al. 2021](https://www.sciencedirect.com/science/article/pii/S266637912100015X) |
|  | YYHKNNKSW | [Tarke et al. 2021](https://www.sciencedirect.com/science/article/pii/S266637912100015X) |
|  | YYVGYLQPRTF | [Tarke et al. 2021](https://www.sciencedirect.com/science/article/pii/S266637912100015X) |
| **B07:02** | **APHGVVFL*** | [Kared et al. 2021](https://pubmed.ncbi.nlm.nih.gov/33427749/) |
|  | EPVLKGVKL | [Tarke et al. 2021](https://www.sciencedirect.com/science/article/pii/S266637912100015X) |
|  | EPVLKGVKL | [Tarke et al. 2021](https://www.sciencedirect.com/science/article/pii/S266637912100015X) |
|  | FPQSAPHGV | [Tarke et al. 2021](https://www.sciencedirect.com/science/article/pii/S266637912100015X) |
|  | IPTNFTISV | [Tarke et al. 2021](https://www.sciencedirect.com/science/article/pii/S266637912100015X) |
|  | KPFERDISTEI | [Tarke et al. 2021](https://www.sciencedirect.com/science/article/pii/S266637912100015X) |
|  | LPFNDGVYF | [Tarke et al. 2021](https://www.sciencedirect.com/science/article/pii/S266637912100015X) |
|  | LPIGINITRF | [Tarke et al. 2021](https://www.sciencedirect.com/science/article/pii/S266637912100015X) |
|  | LPPAYTNSF | [Tarke et al. 2021](https://www.sciencedirect.com/science/article/pii/S266637912100015X) |
|  | LPQGFSAL | [Tarke et al. 2021](https://www.sciencedirect.com/science/article/pii/S266637912100015X) |
|  | MIAQYTSAL | [Saini et al. 2021](https://www.science.org/doi/10.1126/sciimmunol.abf7550) |
|  | QPTESIVRF | [Tarke et al. 2021](https://www.sciencedirect.com/science/article/pii/S266637912100015X) |
|  | QPYRVVVL | [Tarke et al. 2021](https://www.sciencedirect.com/science/article/pii/S266637912100015X) |
|  | QPYRVVVLSF | [Tarke et al. 2021](https://www.sciencedirect.com/science/article/pii/S266637912100015X) |
|  | SPRRARSVA | [Schulien et al. 2021](https://www.nature.com/articles/s41591-020-01143-2) |
|  | SPRRARSVA | [Schulien et al. 2021](https://www.nature.com/articles/s41591-020-01143-2) |
|  | TPCSFGGVSV | [Tarke et al. 2021](https://www.sciencedirect.com/science/article/pii/S266637912100015X) |
|  | TPCSFGGVSV | [Tarke et al. 2021](https://www.sciencedirect.com/science/article/pii/S266637912100015X) |
|  | TPINLVRDL | [Tarke et al. 2021](https://www.sciencedirect.com/science/article/pii/S266637912100015X) |
|  | TPINLVRDL | [Tarke et al. 2021](https://www.sciencedirect.com/science/article/pii/S266637912100015X) |
| **B08:01** | INITRFQTL | [Tarke et al. 2021](https://www.sciencedirect.com/science/article/pii/S266637912100015X) |
|  | KIYSKHTPI | [Tarke et al. 2021](https://www.sciencedirect.com/science/article/pii/S266637912100015X) |
|  | LPQGFSAL | [Tarke et al. 2021](https://www.sciencedirect.com/science/article/pii/S266637912100015X); [Saini et al. 2021](https://www.science.org/doi/10.1126/sciimmunol.abf7550) |
|  | MIAQYTSAL | [Tarke et al. 2021](https://www.sciencedirect.com/science/article/pii/S266637912100015X) |
|  | MIAQYTSAL | [Tarke et al. 2021](https://www.sciencedirect.com/science/article/pii/S266637912100015X) |
|  | NITRFQTL | [Tarke et al. 2021](https://www.sciencedirect.com/science/article/pii/S266637912100015X) |
|  | QPYRVVVL | [Tarke et al. 2021](https://www.sciencedirect.com/science/article/pii/S266637912100015X) |
|  | SIIAYTMSL | [Tarke et al. 2021](https://www.sciencedirect.com/science/article/pii/S266637912100015X) |
|  | SPRRARSV | [Tarke et al. 2021](https://www.sciencedirect.com/science/article/pii/S266637912100015X) |
|  | TLDSKTQSL | [Tarke et al. 2021](https://www.sciencedirect.com/science/article/pii/S266637912100015X) |
|  | YLQPRTFLL | [Tarke et al. 2021](https://www.sciencedirect.com/science/article/pii/S266637912100015X) |
| **B15:01** | CVADYSVLY | [Saini et al. 2021](https://www.science.org/doi/10.1126/sciimmunol.abf7550) |
|  | LVKNKCVNF | [Saini et al. 2021](https://www.science.org/doi/10.1126/sciimmunol.abf7550) |
|  | RLQSLQTY | [Tarke et al. 2021](https://www.sciencedirect.com/science/article/pii/S266637912100015X) |
|  | RVYSTGSNVF | [Tarke et al. 2021](https://www.sciencedirect.com/science/article/pii/S266637912100015X) |
|  | VASQSIIAY | [Saini et al. 2021](https://www.science.org/doi/10.1126/sciimmunol.abf7550) |
| **B35:01** | FAMQMAYRF | [Tarke et al. 2021](https://www.sciencedirect.com/science/article/pii/S266637912100015X) |
|  | FDNPVLPFNDGVYF | [Tarke et al. 2021](https://www.sciencedirect.com/science/article/pii/S266637912100015X) |
|  | FPQSAPHGVVF | [Tarke et al. 2021](https://www.sciencedirect.com/science/article/pii/S266637912100015X) |
|  | FVSNGTHWF | [Tarke et al. 2021](https://www.sciencedirect.com/science/article/pii/S266637912100015X) |
|  | IPFAMQMAY | [Tarke et al. 2021](https://www.sciencedirect.com/science/article/pii/S266637912100015X) |
|  | LGAENSVAY | [Tarke et al. 2021](https://www.sciencedirect.com/science/article/pii/S266637912100015X) |
|  | LPFNDGVYF | [Tarke et al. 2021](https://www.sciencedirect.com/science/article/pii/S266637912100015X) |
|  | LPIGINITRF | [Tarke et al. 2021](https://www.sciencedirect.com/science/article/pii/S266637912100015X) |
|  | LPPAYTNSF | [Tarke et al. 2021](https://www.sciencedirect.com/science/article/pii/S266637912100015X) |
|  | LPPLLTDEM | [Tarke et al. 2021](https://www.sciencedirect.com/science/article/pii/S266637912100015X) |
|  | LTDEMIAQY | [Tarke et al. 2021](https://www.sciencedirect.com/science/article/pii/S266637912100015X) |
|  | NATRFASVY | [Tarke et al. 2021](https://www.sciencedirect.com/science/article/pii/S266637912100015X) |
|  | QIPFAMQMAY | [Tarke et al. 2021](https://www.sciencedirect.com/science/article/pii/S266637912100015X) |
|  | QPTESIVRF | [Tarke et al. 2021](https://www.sciencedirect.com/science/article/pii/S266637912100015X) |
|  | SANNCTFEY | [Tarke et al. 2021](https://www.sciencedirect.com/science/article/pii/S266637912100015X) |
|  | TSNQVAVLY | [Tarke et al. 2021](https://www.sciencedirect.com/science/article/pii/S266637912100015X) |
|  | VASQSIIAY | [Tarke et al. 2021](https://www.sciencedirect.com/science/article/pii/S266637912100015X) |
|  | WTAGAAAYY | [Tarke et al. 2021](https://www.sciencedirect.com/science/article/pii/S266637912100015X) |

* BNT162b2 vaccine-derived immunogenic epitopes identified in this study.

**Supplemental Figures**

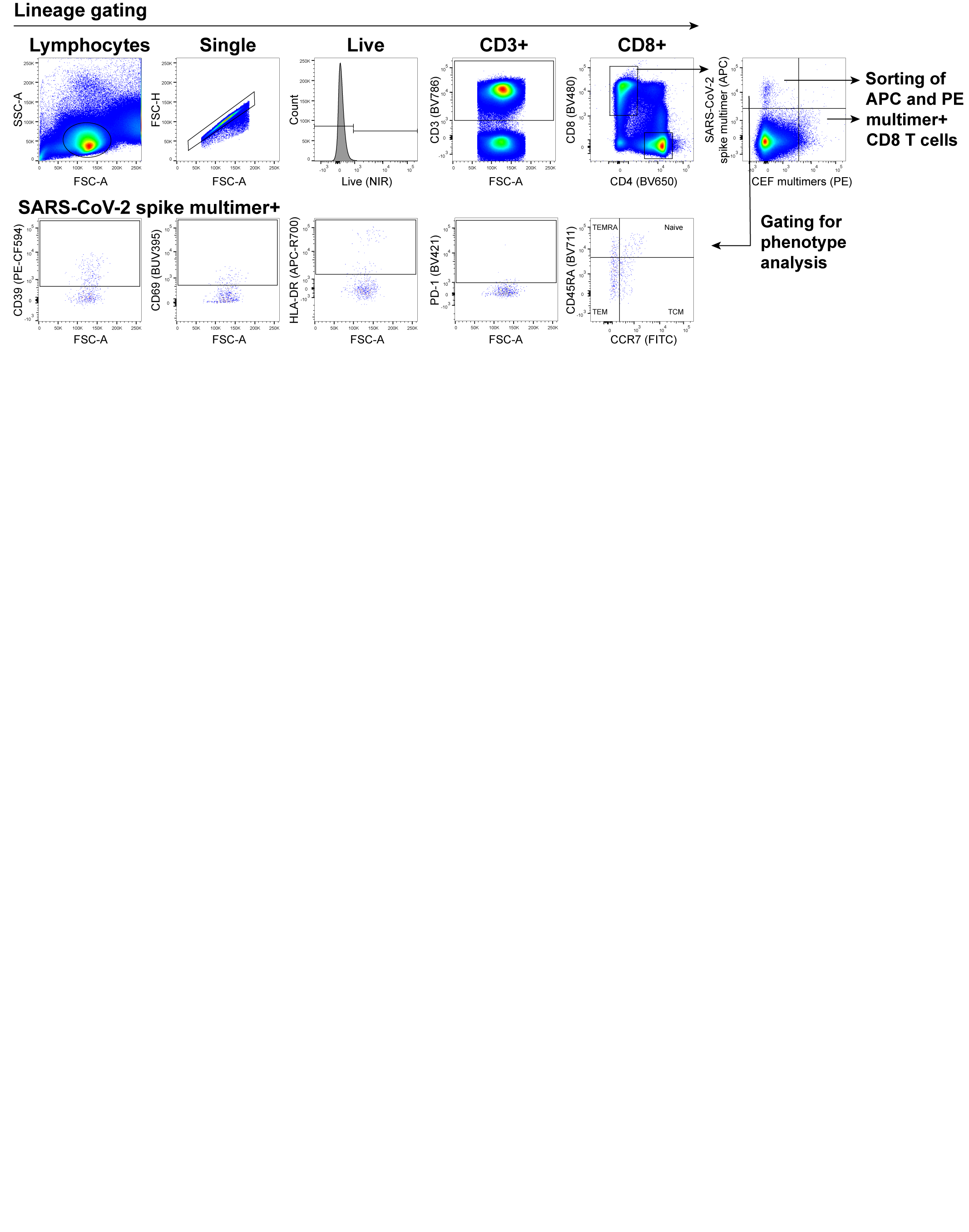

**Supplemental Figure 1. Gating strategy for sorting multimer^+^ CD8^+^ T cells and phenotype analysis.** Representative flow cytometry plots for gating strategy on HM patients and healthy donors PBMCs stained with DNA-barcoded pMHC multimers and surface antibody markers to sort SARS-CoV-2 (APC) and CEF (PE) multimer^+^ CD8^+^ T cells and to quantify multimer^+^ CD8^+^ T cells expressing phenotype markers.

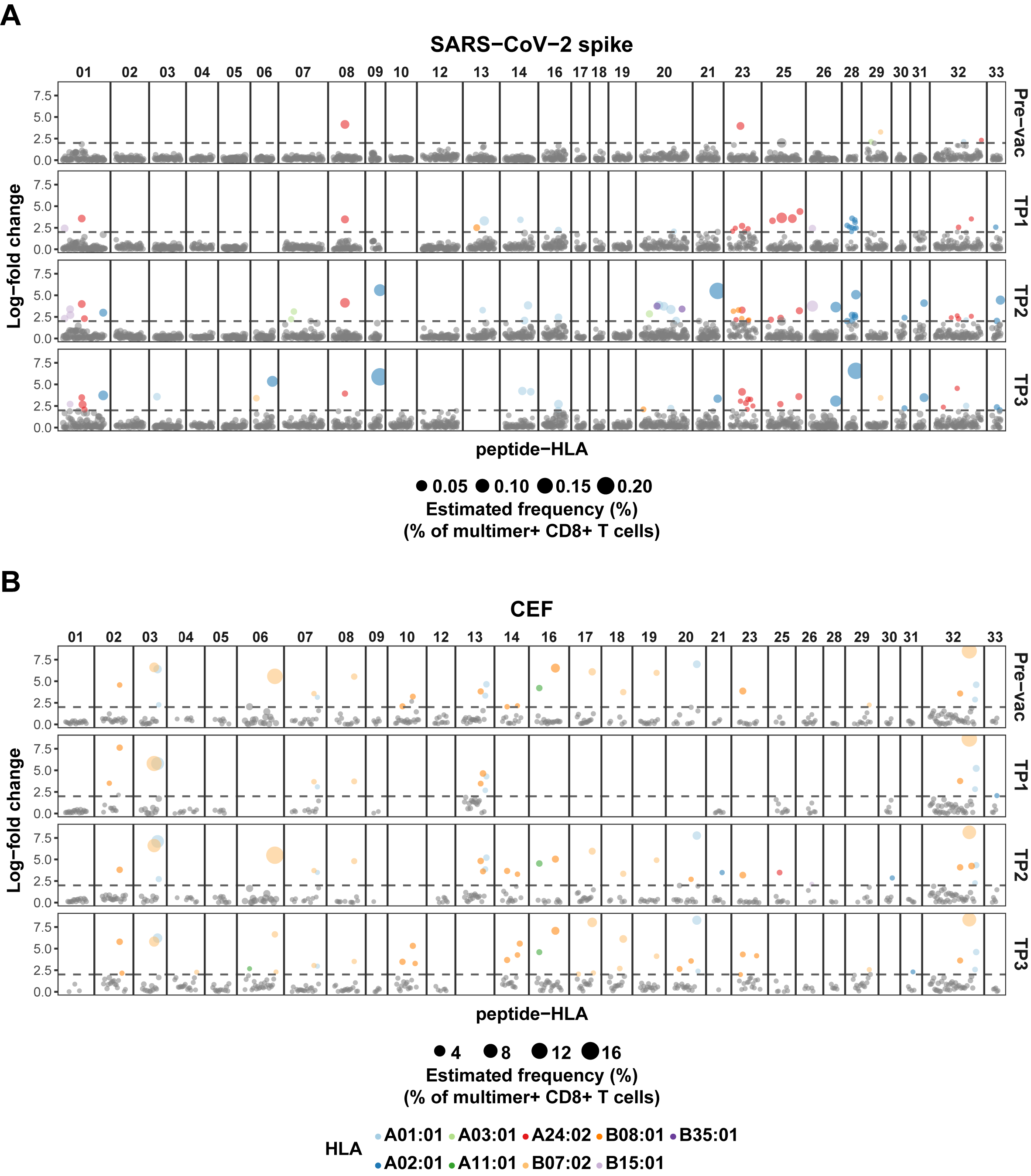

**Supplemental Figure 2. Summary of the CD8^+^ T cell recognition in HM patients.** CD8^+^ T cell recognition to **(A)** SARS-CoV-2 Spike- and **(B)** CEF-derived peptides in each of the HM patients before (Pre-vac) and after vaccination (TP1, TP2 and TP3). Each dot represents one peptide-HLA combination per patients, their size is proportional to the estimated frequency (%) calculated from the percentage read count of the associated barcode out of the percentage of CD8^+^ multimer^+^ T cells, and colored according to their HLA-pair.

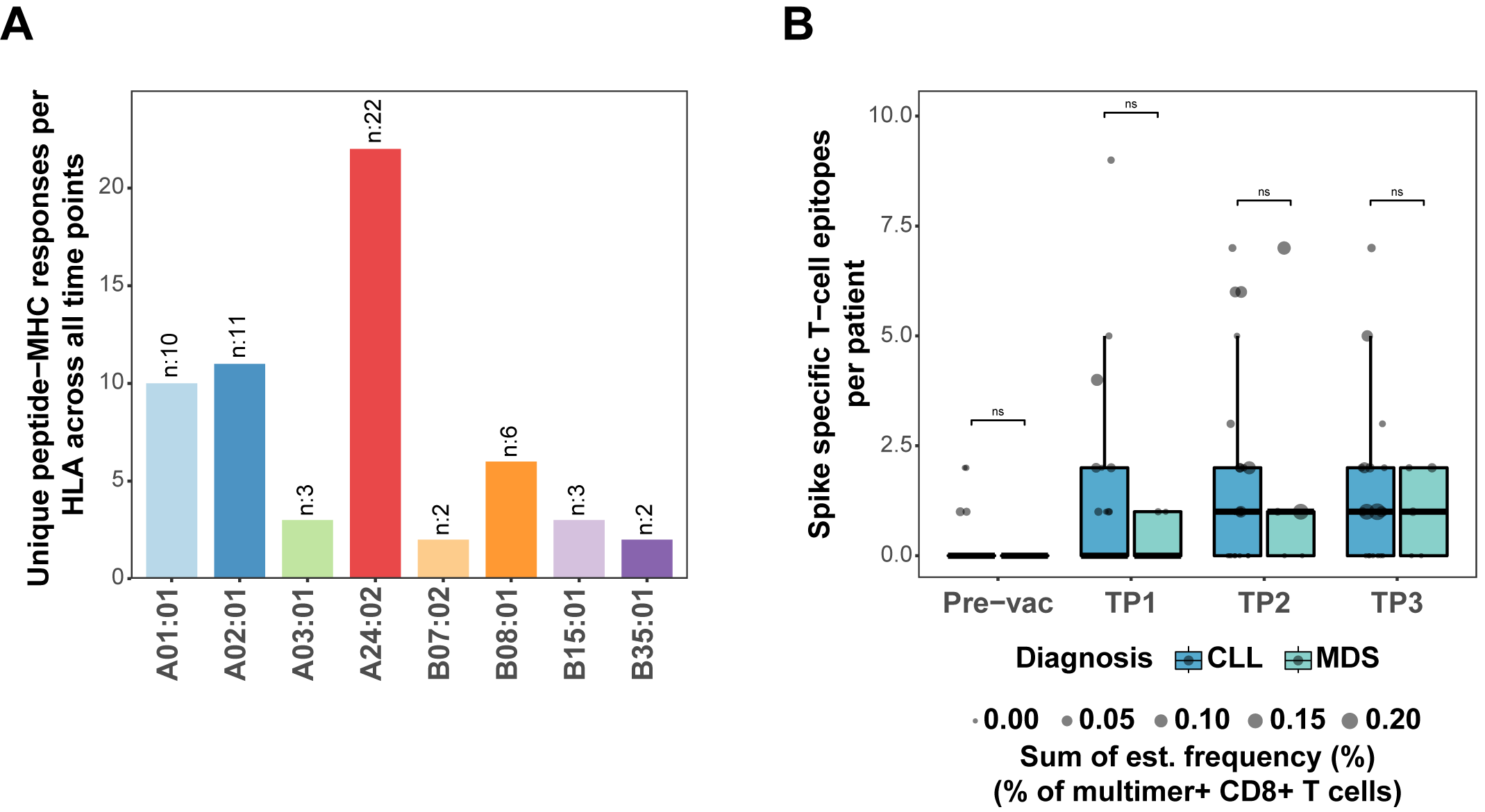

**Supplemental Figure 3. Multimer^+^ CD8^+^ T cell analysis. (A)** Bar plot summarizes the number of unique HLA-specific SARS-CoV-2 Spike epitopes identified in the HM patients across all the time points analyzed. **(B)** Box plot compares the number of SARS-CoV-2 Spike specific T cell epitopes per patient between CLL and MDS diagnosis across the 4 time points. The size of the dots is proportional to the sum of the estimated frequencies (%) of multimer^+^ CD8^+^ T cells for the significant responses in each individual. Mann-Whitney test, Pre-vac CLL vs. MDS (p = 0.324), TP1 CLL vs. MDS (p = 0.475), TP2 CLL vs. MDS (p = 0.771), TP3 CLL vs. MDS (p = 0.845).

.

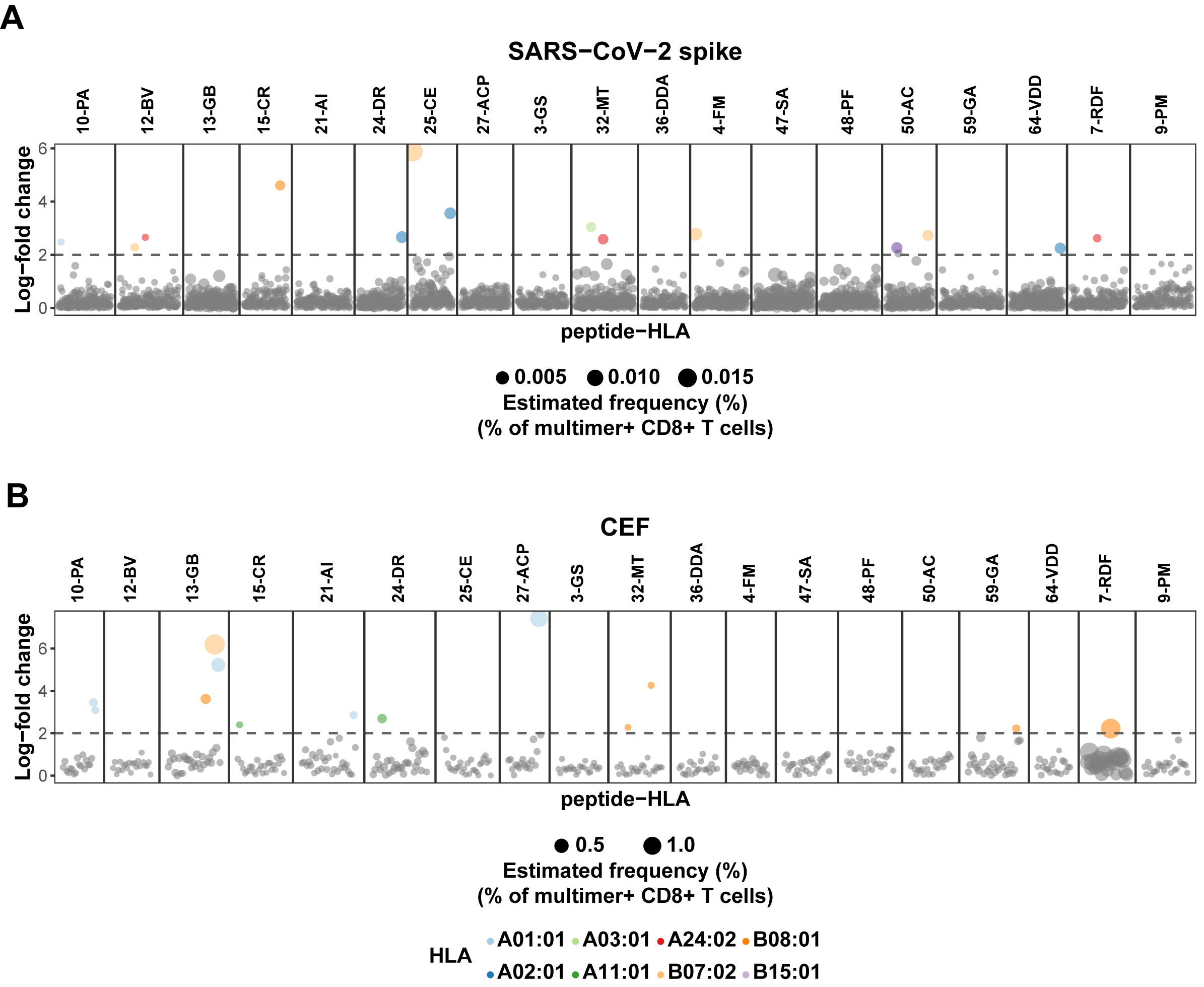

**Supplemental Figure 4. Summary of the CD8^+^ T cell recognition in healthy donors.** T cell recognition to **(A)** SARS-CoV-2 Spike- and **(B)** CEF-derived peptides in each of the health donors before (Pre-vac) and after vaccination (TP1, TP2 and TP3). Each dot represents one peptide-HLA combination per patients, their size is proportional to the estimated frequency (%) calculated from the percentage read count of the associated barcode out of the percentage of CD8^+^ multimer^+^ T cells, and they are colored according to their HLA.

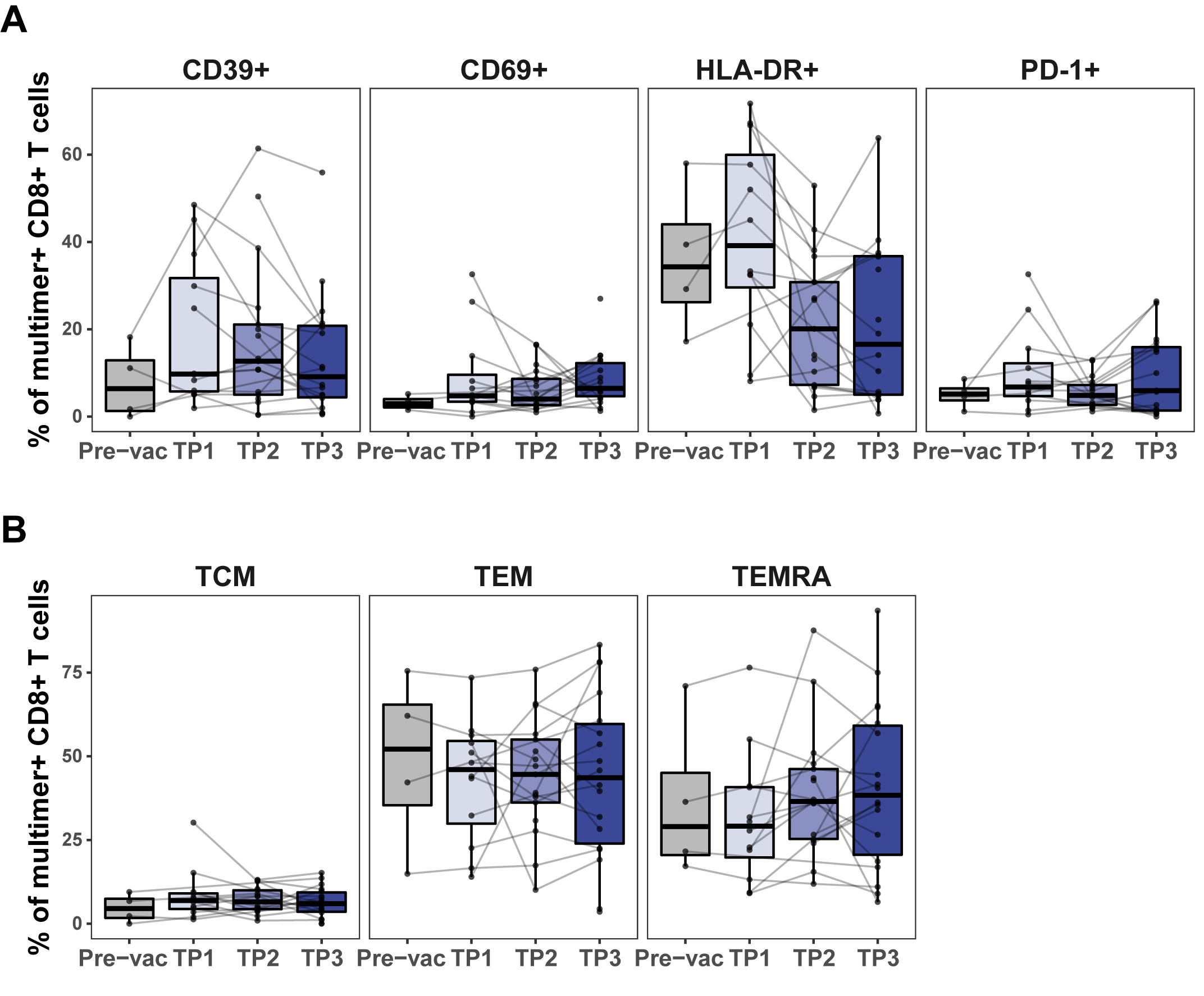

**Supplemental Figure 5. Phenotype analysis for SARS-CoV-2 Spike pMHC multimer^+^ CD8^+^ T cells. (A)** Box plots indicating the percentage of Spike multimer^+^ CD8^+^ T cells expressing the surface markers (CD39, CD69, HLA-DR and PD-1). All p-values > 0.06 using the Kruskal–Wallis one-way ANOVA with Dunn’s correction adjusting p-values with the Bonferroni method. **(B)** The fraction of memory (TCM, TEM and TEMRA) Spike multimer^+^ CD8^+^ T populations based on the expression of CD45RA and CCR7 cell surface markers. Kruskal–Wallis one-way ANOVA with Dunn’s correction adjusting p-values with the Bonferroni method, all p-values > 0.5.

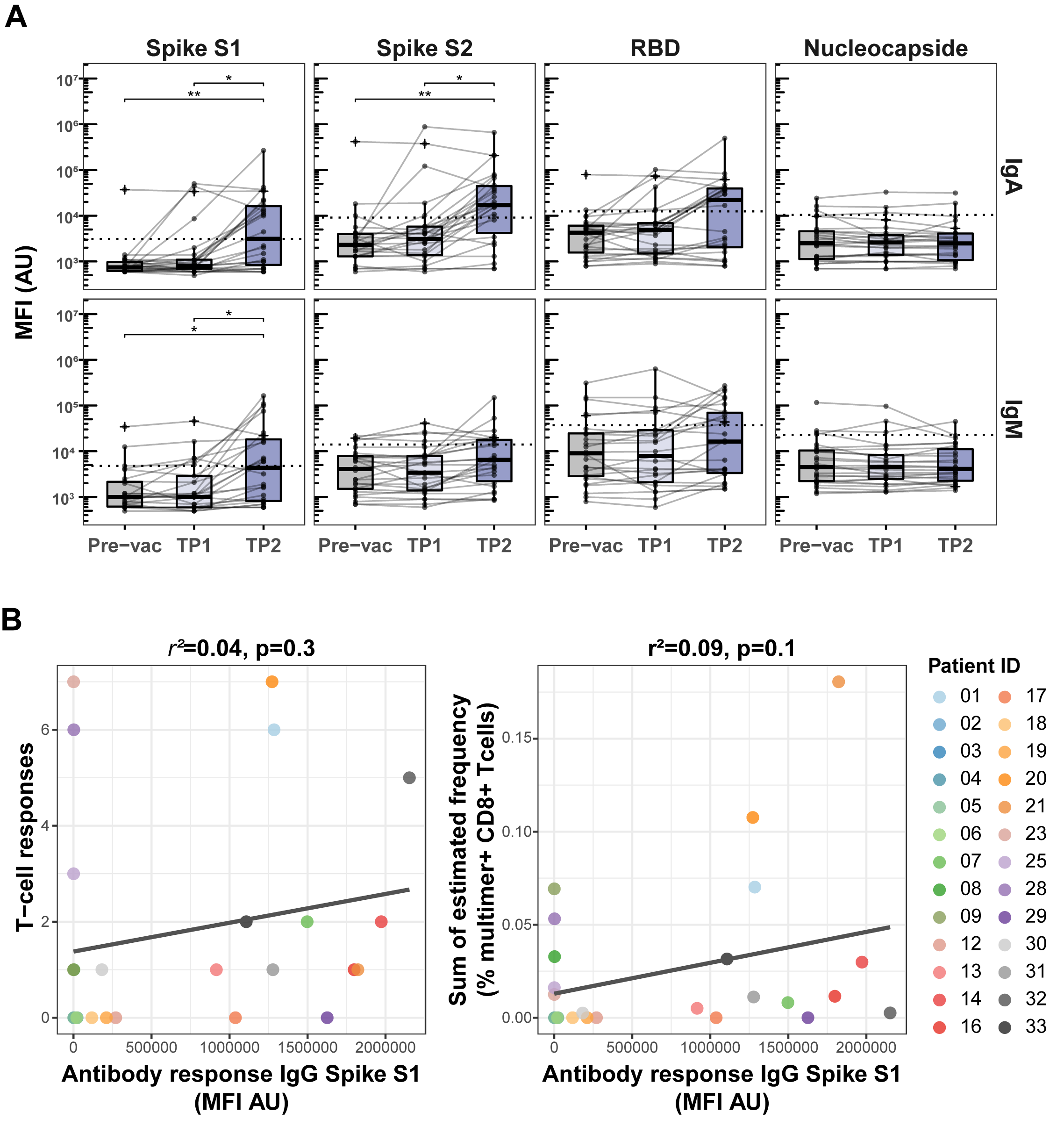

**Supplemental Figure 6. Analysis of antibodies against SARS-CoV-2 antigens. (A)** Box plots showing the levels of IgA and IgM antibodies against SARS-CoV-2 Spike protein subunits S1 and S2, the Spike receptor-binding domain (RBD), and nucleocapside (N) protein in the HM patients at Pre-vac, TP1 and TP2. Threshold for antibody response was set as ≥4-fold increase in geometric mean of the MFI value for pre-vac time point. Kruskal–Wallis one-way ANOVA with Dunn’s correction adjusting p-values with the Bonferroni method, ** (p < 0.01) and * (p ≤ 0.05). **(B)** Plots showing the correlation between the levels of IgG antibody against SARS-CoV-2 Spike protein subunit S1 and **(left)** SARS-CoV-2 Spike specific T cells responses and **(right)** sum of the estimated frequencies (%) for the SARS-CoV-2 Spike specific T cell responses for the HM patients at TP2. The correlation coefficient (r^2^) and p-values are indicated at the top of each plot.

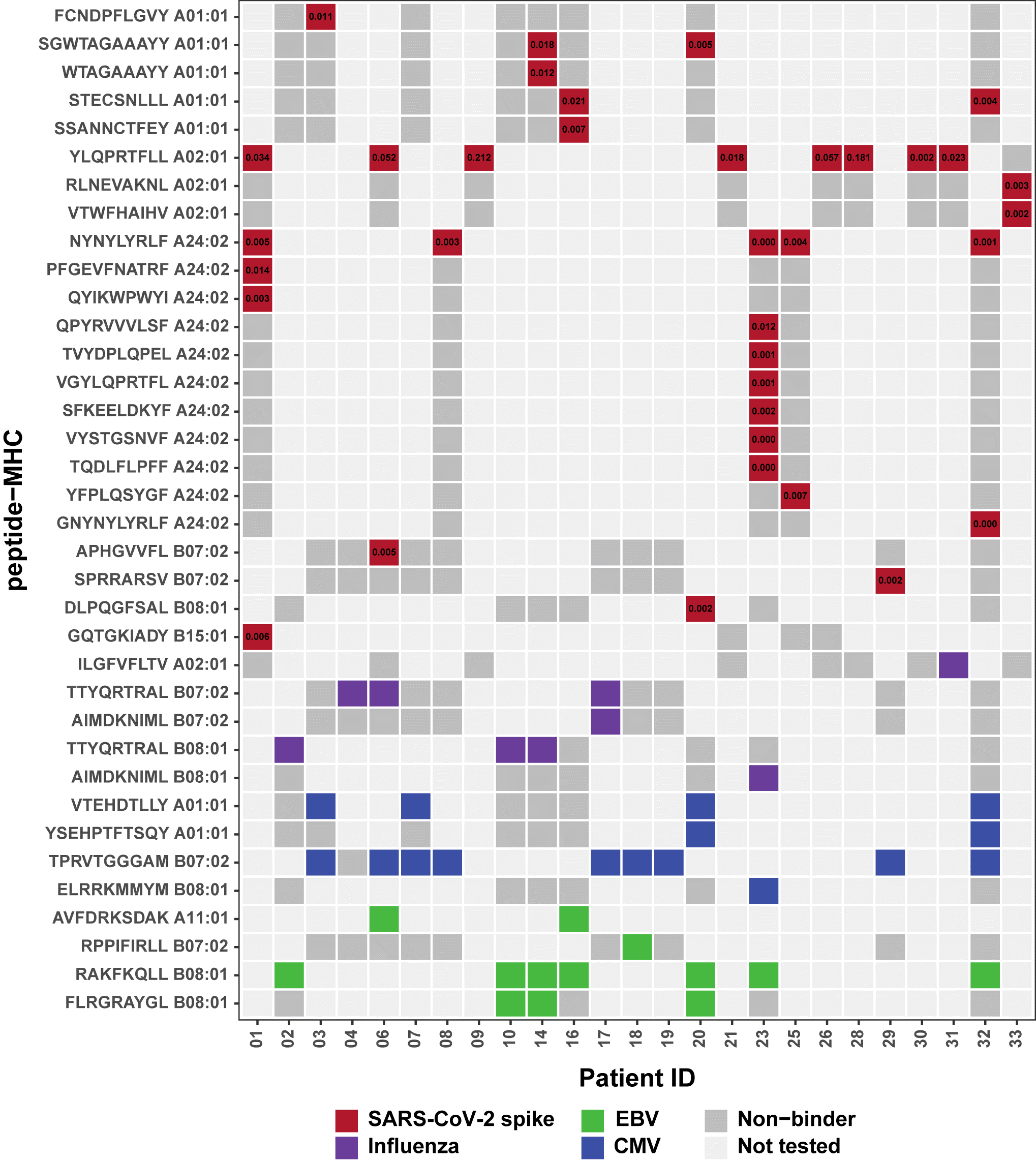

**Supplemental Figure 7. Heatmap plot with patient-specific long-term memory CD8^+^ T cell responses.** Summary of SARS-CoV-2 Spike and CEF-specific epitopes identified in HM patients at TP3. The T cell frequency (%) for the Spike-specific epitopes identified is shown for each HM patient with a positive response.

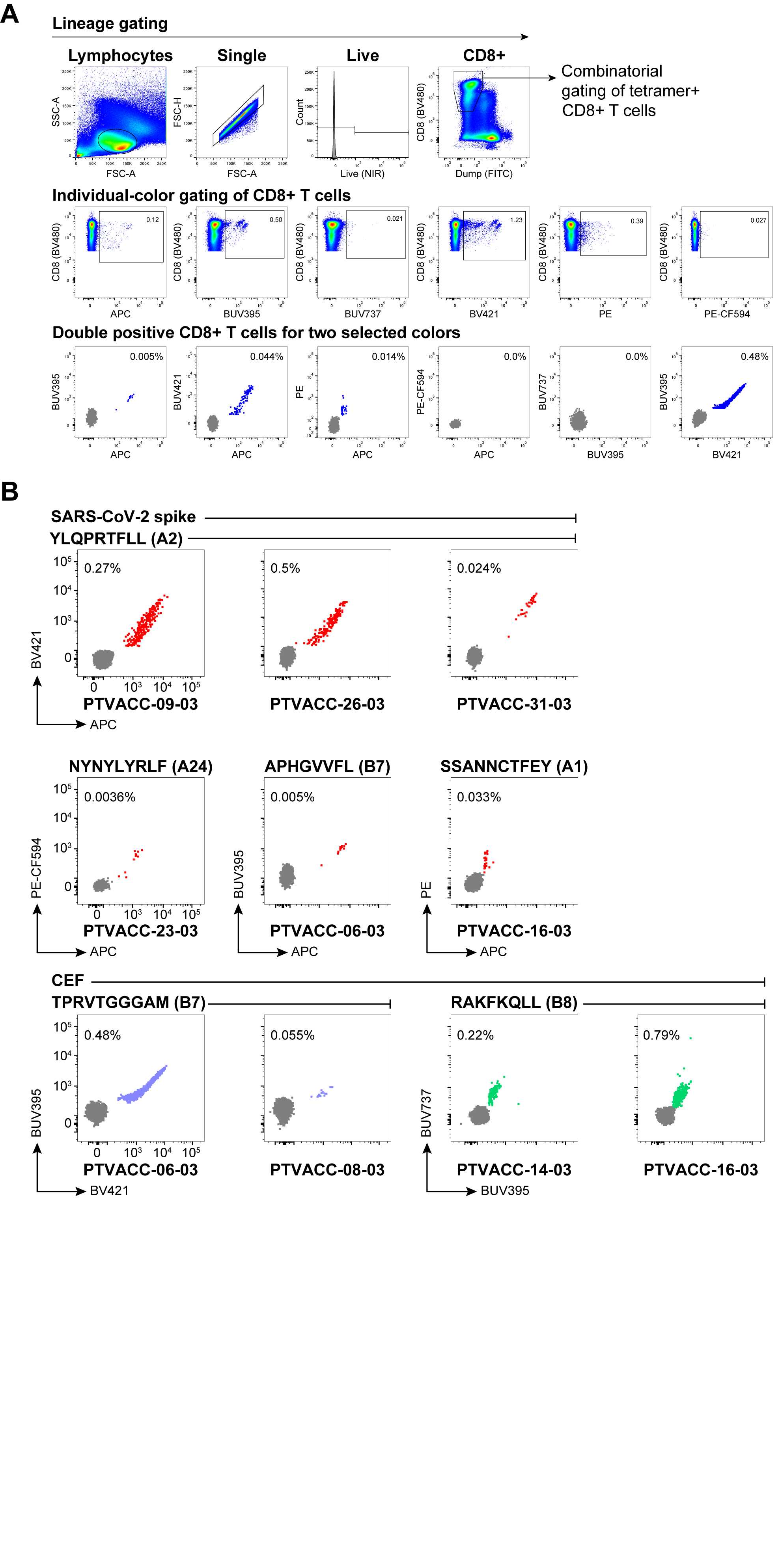

**Supplemental Figure 8. Validation of selected SARS-CoV-2 Spike and CEF epitopes using pMHC tetramers. (A)** Representative flow cytometry plots with gating strategy to identify SARS-CoV-2 Spike- and CEF- specific T cells using combinatorial fluorescently labeled pMHC tetramers. Individual-color gating of CD8^+^ T cells was used to select double positive cells in two tetramer colors and negative in the remaining colors. **(B)** Combinatorial tetramer analysis in HM patients (TP3) of SARS-CoV-2 Spike and CEF derived epitopes identified by DNA barcoded multimers analysis.

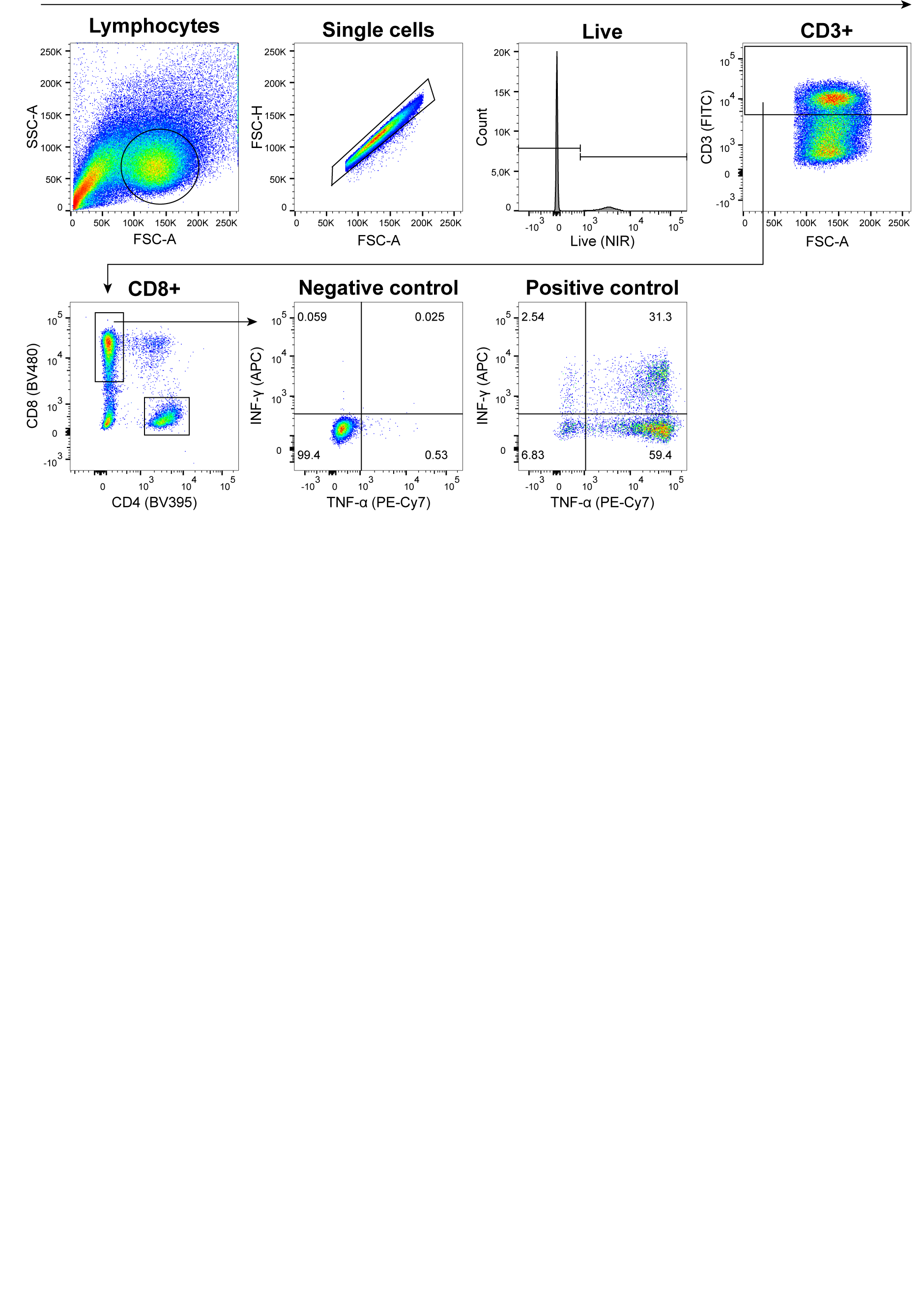

**Supplemental Figure 9. Gating strategy for the functional evaluation of SARS-CoV-2 Spike specific CD8^+^ T cell responses in HM patients.** Representative flow cytometry plots showing the gating strategy to measure intracellular cytokines IFN-γ, and TNF-α of expanded T cells upon stimulation with YLQPRTFLL (HLA-A0201) peptide. Cells incubated with no peptide were used as negative control and cells incubated with leukocyte activation cocktail were used as positive control. The numbers on the plot indicate the frequency (%) of CD8^+^ T cells positive for the analyzed cytokines.
